# Uncovering hidden phylo- and ecogenomic diversity of the widespread methanotrophic genus *Methylobacter*

**DOI:** 10.1101/2025.09.26.678804

**Authors:** Magdalena Wutkowska, Justus A. Nweze, Vojtěch Tláskal, Julius E. Nweze, Anne Daebeler

## Abstract

The globally distributed genus *Methylobacter* plays a crucial role in mitigating methane emissions from diverse ecosystems, including freshwater and marine habitats, wetlands, soils, sediments, groundwater, and landfills. Despite their frequent presence and abundance in these systems, we still know little about the genomic adaptations that they exhibit. Here, we used a collection of 97 genomes and metagenome-assembled genomes to ecogenomically characterise the genus. Our analyses suggest that the genus *Methylobacter* may contain more species than previously thought, with >30 putative species clusters. Some species clusters shared >98.65% sequence identity of the full-length 16S rRNA gene, demonstrating the need for genome-resolved species delineation. The ecogenomic differences between *Methylobacter* spp. include various combinations of methane monooxygenases, multigene loci for alternative dissimilatory metabolisms related to hydrogen, sulphur cycling, and denitrification, as well as other lifestyle-associated functions. Additionally, we describe and tentatively name the two new *Methylobacter* species, which we recently cultured from sediment of a temperate eutrophic fishpond, as *Methylobacter methanoversatilis*, sp. nov. and *Methylobacter spei*, sp. nov. Overall, our study highlights previously unrecognised species diversity within the *Methylobacter* genus, their diverse metabolic potential, versatility, as well as the presence of distinct genomic adaptations for thriving in various environments.

## INTRODUCTION

Methane (CH_4_) is a potent greenhouse gas with an 84-fold greater warming potential than carbon dioxide (CO_2_) over the first 20 years after release (Myhre *et al*. 2013). However, the overall radiative forcing caused by CH_4_ is likely heavily underestimated by approx. 25% (Etminan *et al*. 2016). The CH_4_ level in Earth’s atmosphere has been steadily increasing since the Industrial Revolution. In 2025, the global monthly mean abundance of atmospheric CH_4_ concentration surpassed 1.93 ppm (Lan *et al*. 2022, version 2025-09), nearly tripling the preindustrial 0.7 ppm (Sapart *et al*. 2012). The largest natural sources of CH_4_ are global freshwater systems, including wetlands, which contribute over 50% of total emissions (Jackson *et al*. 2020). Methanotrophs, microorganisms that can use CH_4_ as their only carbon and energy source, can consume >90% of *in situ*-produced CH_4_ before it reaches the atmosphere (King 1990; King, Roslev and Skovgaard 1990; Oremland and Culbertson 1992; Michaud *et al*. 2017). However, it is uncertain whether the efficiency of CH_4_ removal can be sustained in the face of the rapid progression of climate change.

In many ecosystems that produce large amounts of CH_4_, gammaproteobacterial methanotrophs from the genus *Methylobacter* can dominate the active methanotrophic community (Nercessian *et al*. 2005; Tveit *et al*. 2013; Rissanen *et al*. 2018; Smith *et al*. 2018; Savvichev *et al*. 2021; Deng *et al*. 2024; Li *et al*. 2025). Members of the *Methylobacter* genus are widespread, having been identified in samples collected from all continents, predominantly in diverse freshwater, wetland, and soil habitats (Rodrigues *et al*. 2025). They are known to oxidise CH_4_ with the particulate CH_4_ monooxygenase (pMMO), assimilate carbon through the ribulose monophosphate pathway (RuMP), and use ubiquinone Q-8 as a major respiratory lipoquinone (Bowman *et al*. 1993; Bowman 2006). The majority of *Methylobacter* isolates grow relatively quickly and efficiently oxidise CH_4_, even at low temperatures (Tveit *et al*. 2023). In fully oxic conditions, CH_4_ oxidation rates can reach up to 0.60 μmol CH_4_ 10^8^ cells^-1^ hour^-1^ for *M. luteus* at 27°C (Tveit *et al*. 2023). The cultured species grow optimally at temperatures between 23–30°C (Bowman *et al*. 1993; Hanson and Hanson 1996; Bowman 2014; Bodelier *et al*. 2019), with few psychrophilic strains (Omelchenko, Vasilyeva and Zavarzin 1993; Khanongnuch *et al*. 2022; Patil *et al*. 2024), widespread psychrotolerance (Wartiainen *et al*. 2006; Roldán and Menes 2023), and only some thermotolerant species with optima >35°C (Lidstrom 1988). *Methylobacter* spp., although geographically widely distributed, are predominantly found in non-acidic pH ranges (Seppey *et al*. 2023), with a few exceptions (Nguyen *et al*. 2018; Hogendoorn *et al*. 2021; Nweze *et al*. 2024).

The genus *Methylobacter* was initially named in 1970 (Whittenbury, Davies and Davey 1970; Whittenbury, Phillips and Wilkinson 1970). However, the name was only officially (taxonomically) introduced two decades later, reclassifying known methanotrophs based on the DNA-DNA hybridisation values and phospholipid fatty acid compositions (Bowman *et al*. 1993). A genome-based reclassification of methanotrophs, including the genus *Methylobacter*, corrected some misclassifications, which had previously contributed to a polyphyletic character of the genus, and identified 10 species-level clusters in the genus *Methylobacter* (Orata *et al*. 2018).

In this study, we used 97 genomes and MAGs classified as *Methylobacter*, retrieved in May 2024 from the NCBI Genome portal, to comprehensively characterise them regarding their potential metabolic capabilities in the context of phylogenomics. A special focus was directed towards the presence and genomic organisation of different CH_4_ monooxygenase forms, dissimilatory metabolisms other than CH_4_ oxidation, and genomic adaptations to O_2_-depleted conditions. Additionally, we characterise two recently obtained novel *Methylobacter* spp. (Wutkowska and Daebeler 2024), which we tentatively name *Methylobacter spei*, sp. nov., and *Methylobacter methanoversatilis*, sp. nov.

## MATERIALS AND METHODS

### *Methylobacter* genomes and MAGs

On May 21, 2024, we retrieved all 97 available genomes and MAGs classified as *Methylobacter* from the NCBI Genome database (Table S1, Fig. 1). Genomes of isolates included: *M. luteus* 98 or IMV-B-3098 previously *M. bovis* (Whittenbury, Davies and Davey 1970; Whittenbury, Phillips and Wilkinson 1970)*; M. whittenburyi* (formerly *M. capsulatus* UCM-B-3033) (Whittenbury, Phillips and Wilkinson 1970; Hamilton *et al*. 2015); *M. marinus* A45 (formerly *Methylomonas methanica* A4) (Lidstrom 1988; Flynn *et al*. 2016); *M. tundripaludum* SV96 (Wartiainen *et al*. 2006; Svenning *et al*. 2011); *M. psychrophilus* Z-0021 (Omelchenko, Vasilyeva and Zavarzin 1993; Omelchenko *et al*. 1996b; Rissanen *et al*. 2022); *M.* sp. strain BBA5.1 (Smith, Costello and Lidstrom 1997; Flynn *et al*. 2016); *M. sp.* 21/22 and 31/32 (Beck *et al*. 2013; Kalyuzhnaya *et al*. 2015; Oshkin *et al*. 2015); ‘*Ca.* M. oryzae’ KRF1 (Khatri *et al*. 2020); ‘*Ca*. M. coli’ BlB1 (Khatri *et al*. 2021); *M*. sp. YRD-M1 (Hao *et al*. 2022), *M*. sp. Wu8 (Wutkowska and Daebeler 2024), and *M. svalbardiensis* LS7-T4A (Patil *et al*. 2024). A recent reclassification proposed *M. whittenbury* and *M. marinus* to be the heterotypic synonyms, as their genomes share a very high degree of similarity (89% dDDH and 99% ANI) (Orata *et al*. 2018), with *M. marinus* being the recommended valid name, therefore we did not include a *M. whittenbury* genome in our analyses.

**Figure 1.**
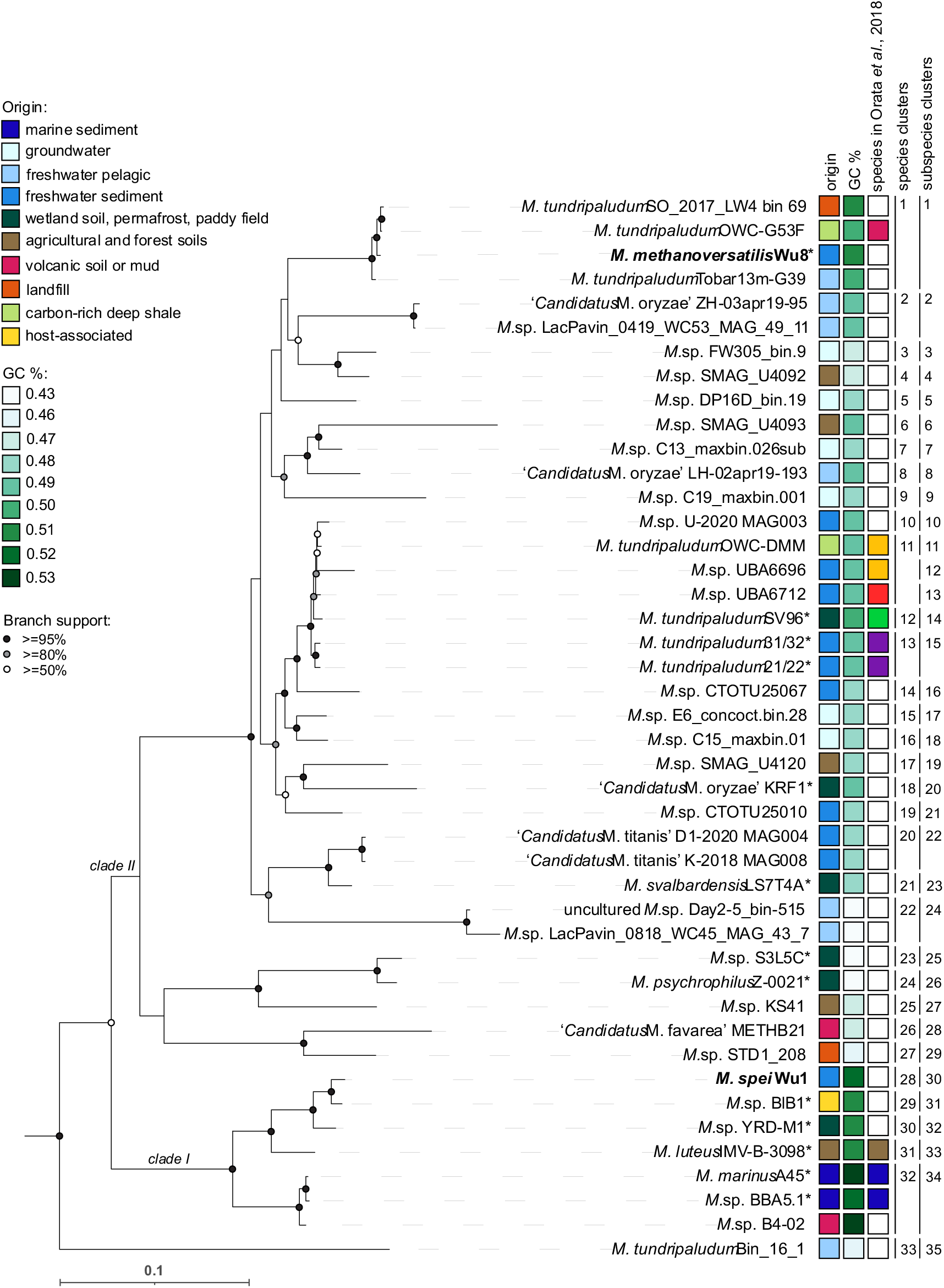
Phylogeny of the genus *Methylobacter* based on concatenated amino acid sequences of 71 single-copy genes from 44 high-quality non-redundant *Methylobacter* spp. genomes and MAGs. Genomes from axenic cultures are indicated by asterisks (*), and genomes from our recently cultured strains are shown in **bold**. Coloured squares indicate the origin habitat type, GC content, and species included and delineated in Orata *et al*. (2018) (each colour represents a distinct species). Numbers and vertical lines indicate species and subspecies inferred by TYGS. The genome of *Methylobacterium lacus* LW14 (Kalyuzhnaya *et al*. 2015) was used as an outgroup. Branch bootstrap support is indicated on the nodes as black (≥95%), grey (≥80%), or white (>50%) circles. The scale bar indicates 0.1 amino acid substitutions per site.

*Methylobacter* spp. that have been sequenced from enrichment cultures belong to: *M*. sp. KS41 (Nguyen *et al*. 2018); ‘*Ca.* M. titanis’ sp. nov. strains K-2018 and D1-2020, as well as *M*. sp. U-2020 MAG003 (Roldán and Menes 2023), and *M*. sp. strains Wu1 (Wutkowska and Daebeler 2024). The remaining MAGs originated from environmental samples (Parks *et al*. 2017; Woodcroft *et al*. 2018; Pedron *et al*. 2019; Zheng *et al*. 2020; Buck *et al*. 2021; Hogendoorn *et al*. 2021; Rissanen *et al*. 2021; Magnuson *et al*. 2022; Cabello-Yeves *et al*. 2023; Grégoire, George and Hug 2023; Jaffe *et al*. 2023; Ma *et al*. 2023; Ruff *et al*. 2023; Slobodkin *et al*. 2023).

The gene coding sequences were predicted and translated to amino acid sequences using Prodigal v.2.6.3 (Hyatt *et al*. 2010). To assess similarity among the 97 genomes, we calculated average nucleotide identity inferred with *blastp* (ANIb) using pyANI (Pritchard *et al*. 2016) in-built in a pangenome workflow in anvio v.7.1 ‘Hope’ (Eren *et al*. 2021) with third party software DIAMOND (Buchfink, Reuter and Drost 2021) and MUSCLE (Edgar 2004). To reduce redundancy in our dataset, genomes and MAGs with >99% ANIb similarity were grouped, and a representative genome for each group was chosen based on the highest completeness and lowest contamination calculated with CheckM2 v.1.0.1 (detailed summary available in Table S1) (Chklovski *et al*. 2023). Those genomes with >90% completeness and <5% contamination was categorised as ‘high-quality’, although they sometimes consist of many contigs. Subsequently, only representative high-quality genomes and MAGs (n = 44) were retained for all downstream analyses (Table S1, Fig. 1).

### Phylogenomic analysis, genus and species delineation

The phylogenomic analysis was performed using Anvi’o in the Anvi’o development environment (anvio-dev) (Eren *et al*. 2021). The FASTA files containing genome sequences were reformatted to meet Anvi’o’s contigs database requirements using the anvi-script-reformat-fasta command. For each reformatted FASTA file, an Anvi’o contigs database was generated using the anvi-gen-contigs-database command, which used Prodigal (v2.6.3) (Hyatt *et al*. 2010) to identify open reading frames in the DNA sequences and predict protein-coding genes. Conserved genes, including single-copy core genes (SCGs), were annotated and identified using hidden Markov models (HMMs) with the anvi-run-hmms command and the Bacteria_71 HMM profile. The sequences of the best hit for each SCG were extracted and concatenated into a single FASTA file using the anvi-get-sequences-for-hmm-hits command. The concatenated protein sequences were aligned using MAFFT (v7.520) (Katoh and Standley 2013) with the FFT-NS-2 strategy, which was automatically selected using the --auto option. The aligned sequences were then used to construct a maximum-likelihood phylogenetic tree using IQ-TREE (v2.1.1) (Nguyen *et al*. 2015; Minh *et al*. 2020). The IQ-TREE used ModelFinder Plus (Kalyaanamoorthy *et al*. 2017) to automatically select the best-fit substitution model JJT+F+R5, and 1,000 ultrafast bootstrap replicates with UFBoot2 (Hoang *et al*. 2018) to assess branch support.

To determine whether the investigated *Methylobacter* genomes belong to the same genus, we calculated the average amino acid identity (AAI) using the EzAAI toolkit v1.2.2 (Kim, Park and Chun 2021), which extracts and estimates the pairwise similarity using Prodigal (Hyatt *et al*. 2010) and MMseqs2 (Steinegger and Söding 2017). *Methylobacter* species were delineated using the Type-Strain-Genome-Server (TYGS) (Meier-Kolthoff and Göker 2019; Meier-Kolthoff *et al*. 2022). This tool combines several indices and other types of evidence commonly used to distinguish species, including a phylogenomic tree calculated using genome BLAST distance phylogeny (GBDP), digital DNA–DNA hybridisation (dDDH), average nucleotide identity, and differences in genomic GC content. The dDDH scores were obtained using the Genome-to-Genome Distance Calculator 3.0, based on BLAST+ (Camacho *et al*. 2009) with the recommended formula 2 (d_4_) (Meier-Kolthoff *et al*. 2013, 2022), as well as the differences between genomic GC content (Meier-Kolthoff, Klenk and Göker 2014). Species were ultimately delineated with the agreement of all the used metrics, with dDDH being the conclusive one. Two genomes were clustered into the same species when the dDDH value >70% (Wayne *et al*. 1987; Goris *et al*. 2007), and into the same subspecies with >79% (Meier-Kolthoff *et al*. 2014; Meier-Kolthoff and Göker 2019).

We constructed phylogenetic trees with the 16S rRNA gene using nucleotide sequences and with protein-coding genes of interest using amino acid sequences (i.e., CH_4_ monooxygenases, ATP synthase subunit c, ATP synthases). In each case, sequences were aligned using MAFFT v7.526 with the --auto option to optimise the alignment process (Katoh and Standley 2013). The branch support in protein phylogenetic trees was obtained with the ultrafast bootstrap method with 1,000 replicates (Hoang *et al*. 2018) implemented in the IQ-TREE 2 (Minh *et al*. 2020). Trees were visualised in iTOL v7 (Letunic and Bork 2024). Visualisation of the genomic region of soluble CH_4_ monooxygenase was performed using the gggenomes package (Hackl *et al*. 2024) in R v4.0 (R Core Team 2024).

### Annotations of open reading frames

We screened all representative high-quality *Methylobacter* genomes and MAGs for the presence of genes encoding three CH_4_ monooxygenases and a gas vesicle protein with in-house Hidden Markov Models (HMMs) with the hmmscan/hmmsearch function in HMMER 3.4 (Eddy 2011). The HMM for the major gas vesicle structural protein (encoded by the *gvpA* gene) was constructed from 32 reviewed and aligned amino acid sequences found in the UniProt Knowledgebase (The UniProt Consortium *et al*. 2025). All HMMER hits with a score <120 and an e-value <1e^-30^ were considered as *gvpA*. Additionally, we screened the genomes and MAGs for metabolic genes (i.e., [NiFe] hydrogenases, *soxB*, *narGHI*) using DIAMOND v2.1.9.163 (Buchfink, Xie and Huson 2015) against the collections of reference amino acid sequences (Leung and Greening 2021). Sequences identified as [NiFe]-hydrogenases were subsequently classified into specific groups using HydDB (Søndergaard, Pedersen and Greening 2016). All MAGs were also annotated with the automatic pipeline DRAM (Shaffer *et al*. 2020).

To contextualise the distribution of soluble methane monooxygenase (i.e., *mmoX)* and gas protein vesicles (i.e., *gvpA)* among gammaproteobacterial methanotrophs, we downloaded 1,067 available genomes and MAGs listed as belonging to the order *Methylococcales* from NCBI on August 12, 2024, and used in-house HMMs to identify the presence of these genes.

Biosynthetic gene clusters were identified in the high-quality non-redundant genomes and MAGs with the Minimum Information about a Biosynthetic Gene cluster database v4.0 (Zdouc *et al*. 2025) using blastp in DIAMOND v2.1.9.163 (Buchfink, Xie and Huson 2015), and antiSMASH v7.1.0 (Blin *et al*. 2023), both with stringent parameters.

### Predicting optimal growth conditions

Since the majority of *Methylobacter* species have not been cultured yet, little is known about their ecological preferences and optimal growth conditions. Therefore, we identified putative oxygen preferences, optimal temperature, salinity, and pH levels based on genomic amino acid compositions of all high-quality genomes and MAGs using GenomeSPOT with default statistical models (Barnum *et al*. 2024). The obtained predictions were compared with available data from cultured *Methylobacter* spp.

## RESULTS AND DISCUSSION

### Phylogenomic analysis and genome characteristics of *Methylobacter*

The *Methylobacter* genomes and MAGs differed substantially in general characteristics, such as size (3,452,370–5,467,791 bp), number of predicted proteins (3,206–5,043), and GC content (44–55%) (Table S1, Fig. 1). The highest GC content occurred in the marine cluster and an adjacent cluster of *Methylobacter* spp. associated with eutrophic habitats and (putative) animal hosts (Table S1, Fig. 1). Accordingly, genomes that belong to psychrophilic strains and species from potentially nutrient-depleted habitats had the lowest GC content among all *Methylobacter* spp., reflecting adaptation to temperature and resource availability as described previously (Foerstner *et al*. 2005; Hu *et al*. 2022; Chuckran *et al*. 2023).

The *Methylobacter* genus contains two recognised, phylogenetic lineages, termed clade I, encompassing species such as *M. luteus* and *M. marinus*, and clade II, which includes the majority of *Methylobacter* spp. including *M. tundripaludum* (Smith *et al*. 2018a). General AAI similarity recommended for clustering genomes/MAGs in one genus ranges between 65–95% (Konstantinidis, Rosselló-Móra and Amann 2017); however, a *Methylobacter*-specific threshold has been set at 74% (Orata *et al*. 2018). The MAG *M. tundripaludum* Bin_16_1, which was the earliest diverging taxon, branched out most deeply in the phylogenomic tree (Fig. 1), shared only 72% AAI with all the genomes/MAGs in clade I, and 73–74% AAI with the remaining genomes/MAGs, including its closest related *Methylobacter* genome, ‘*Ca.* Methylobacter favarea’ (Fig. S1). Moreover, *M. tundripaludum* Bin_16_1 has been classified as belonging to the genus *Crenothrix* in the GTDB Release R226 (Parks *et al*. 2022) (Table S1). According to another measure of genome similarity allowing for genus delineation—a fixed percentage of conserved proteins (POCP) set to 50%, clade II may actually consist of at least five separate genera (Orata *et al*. 2018), whereas the GTDB classification divides clade II into three genera: Methylobacter_A, Methylobacter_B, and Methylobacter_C in the GTDB Release 226 (Parks *et al*. 2022). However, ANIb values suggest that all the analysed genomes and MAGs, including *M. tundripaludum* Bin_16_1, belong to the genus *Methylobacter* based on the >73% ANI similarity threshold (Barco *et al*. 2020). Hence, our analyses suggest that the taxonomy of *Methylobacter*, and possibly that of *Crenothrix*, may need re-evaluation, which is, however, beyond the scope of this study. Our phylogenomic analysis confirmed the separation of the two *Methylobacter* clades (Fig. 1). Additionally, we identified 33 unique species-level clusters based on several methods of species delineation, such as the TYGS clustering (Meier-Kolthoff and Göker 2019; Meier-Kolthoff *et al*. 2022), <95% of ANIb (Jain *et al*. 2018), and dDDH scores of <70 (Goris *et al*. 2007). Our analyses suggest that the *Methylobacter* genus may be composed of more species than previously recognised, with even more species to be identified in the near future through continued sequencing and culturing efforts.

We recently cultured two new *Methylobacter* spp., *M.* sp. Wu1 and *M.* sp. Wu8, from the sediment of a temperate eutrophic fishpond (Wutkowska and Daebeler 2024). The Wu1 strain shared >95% ANIb identity with ‘*Ca.* M. coli BlB1’ (Fig. S4), which was backed up by close clustering in the phylogenomic tree (Fig. 1). However, further evidence, i.e., a dDDH score of 64.3 in a pairwise genome comparison, suggested that they nevertheless likely belong to different species. Interestingly, ‘*Ca.* M. coli BlB1’ has been isolated from the faeces of a herbivore (Khatri *et al*. 2021), and it therefore seems plausible that *M.* sp. Wu1 may in fact originate from cattle manure, which is routinely used for fertilising fishponds in the area (Potužák, Hůda and Pechar 2007) and not from the sediment. Here, we tentatively propose the name *Methylobacter spei*, sp. nov. The genome of the Wu8 strain phylogenomically clustered most closely with three other MAGs of uncultured *Methylobacter* (Fig. 1). These were obtained from various aquatic environments in the northern hemisphere, and one of them, termed *M. tundripaludum* OWC-G53F, has previously been proposed to belong to a new species within the genus (Orata *et al*. 2018). Our ANIb and TYGS analyses corroborated the phylogenetic affiliation of the Wu8 strain with the uncharacterised species-level cluster distantly related to the next cultured relative (Fig. 1) and further suggested that it represents the first cultured member of a novel species for which we propose the name *Methylobacter methanoversatilis*, sp. nov. (Fig. 1; Fig. S4).

The phylogeny of the 18 available full-length 16S rRNA gene sequences from the analysed *Methylobacter* genomes was in congruence with the phylogenomic topology, preserving the major division into two clades (Fig. S2a). However, several distinct *Methylobacter spp.* displayed >99% nucleotide identity for the entire 16S rRNA gene (Khanongnuch *et al*. 2022; Rissanen *et al*. 2022), which is above the recommended 98.65% threshold to delineate bacterial species for this marker (Kim *et al*. 2014). For instance, the identity of the 16S rRNA genes of *M. psychrophilus* Z-0021 and *M. tundripaludum* SV96 is above this threshold, but their genome comparison yielded values below the 95% ANIb and 70% dDDH scores, which are recommended for species delineation (Goris *et al*. 2007; Jain *et al*. 2018), suggesting they are two distinct species. These discrepancies between the 16S rRNA gene-based and the genome-based species delineation can lead to erroneous conclusions when (often <250 bp) sequences of 16S rRNA generated in amplicon studies are used to study *Methylobacter* sp. community composition at the species level. Upon closer inspection, we also noticed that the most commonly targeted region of the 16S rRNA gene, the V4 region, which is also included in the Earth Microbiome Project (Caporaso *et al*. 2018), is highly conserved among several *Methylobacter* species. This was, for example, true for *M. tundripaludum* SV96 and *M. methanoversatilis* Wu8, which differed by only 1 nucleotide in the V4, but by 14 nucleotides in the V7–V8 region (Fig. S3). We therefore conclude that attempting to distinguish between *Methylobacter* species by comparing 16S rRNA gene sequences will often lead to an underestimation of diversity, as has been recently shown for other genera (Alleman *et al*. 2025). The PmoA-based trees (Fig. S2b, Fig. S2c) present a less congruent picture than the 16S rRNA and phylogenomic tree (Fig. S2a and Fig. 1, respectively), which is expected and has been previously shown to yield *Methylobacter* spp. as a polyphyletic genus (Knief 2015; Orata *et al*. 2018).

### A varied repertoire of CH_4_ and methanol oxidation

Nearly all the investigated *Methylobacter* genomes (91%) contained at least one gene cluster coding for particulate CH_4_ monooxygenase, either in its canonical *pmoCAB* (pMMO) or in the sequence-divergent *pxmABC* (pXMO) form (Tavormina *et al*. 2011) (Fig. 2). Almost half of the genomes (45%) encoded for both forms. Additionally, eight genomes (18%) contained the complete inventory for the cytoplasmic, soluble CH_4_ monooxygenase (sMMO) with structural (*mmoXYBZ(D)C*) and regulatory genes, such as *mmoG* (Fig. 2, Fig. 3). A phylogenetic analysis of the protein sequence of the hydrolase alpha subunit of sMMO, MmoX, showed a clear separation of *Methylobacter* MmoX into two clusters. MmoX sequences from *M. psychrophilus* Z-0021 and *M.* sp. S3L5C are affiliated closely with sequences from *Crenothrix polyspora* and *Methylovulum miyakonense* HT12, whereas the other cluster with the remaining MmoX, including sequences from *M. methanoversatilis*, formed a deeper branching, isolated group (Fig. 3B). The latter cluster lost the additional gene coding for an unknown hypothetical protein between *mmoC* and *mmoG*, similarly to *Methylovulum miyakonense* HT12 (Iguchi, Yurimoto and Sakai 2010) (Fig. 3A, Fig. 3B).

**Figure 2.**
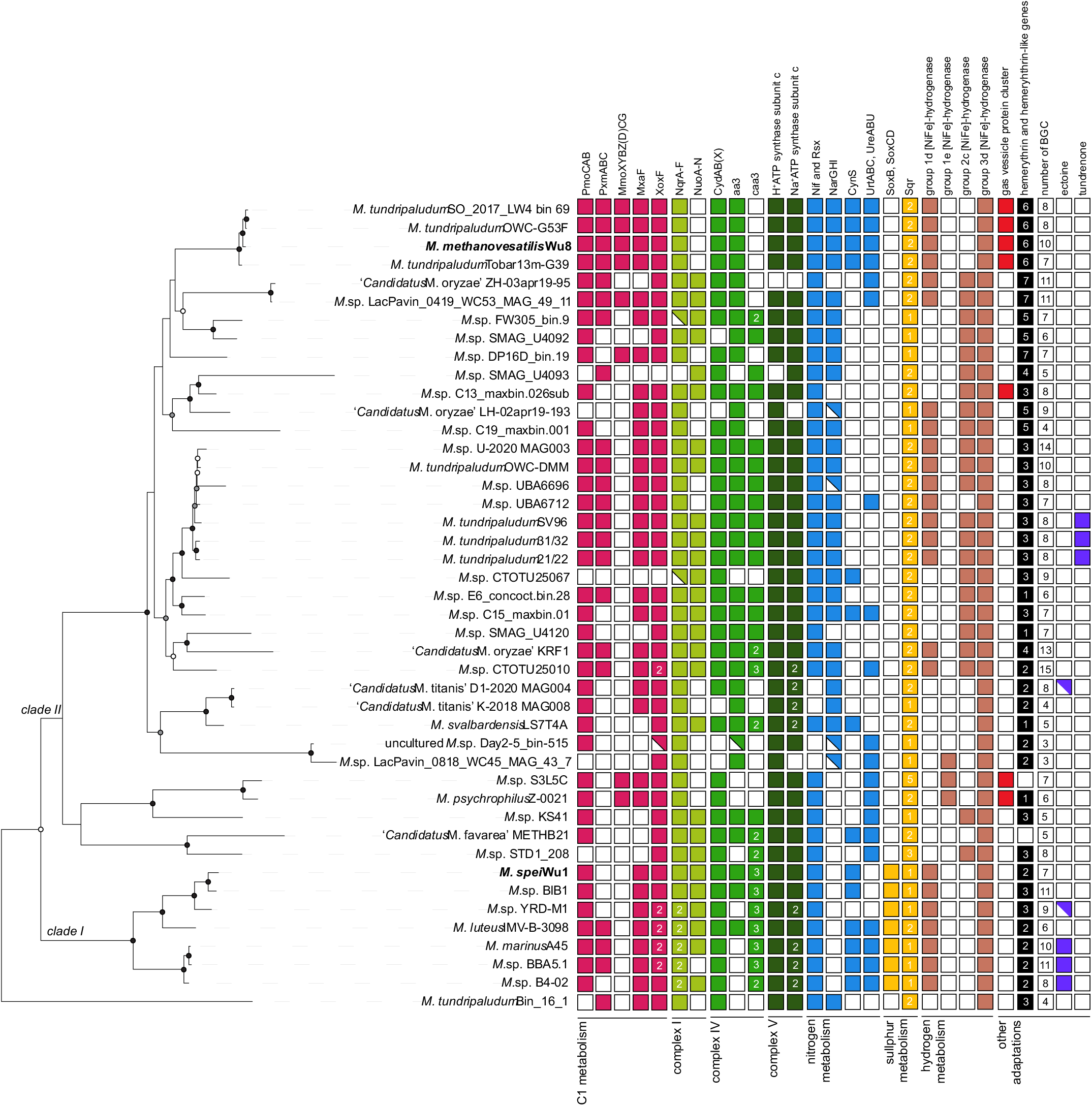
Distribution of distinct metabolic traits in *Methylobacter*. Protein homology was inferred based on the HMM models, alignment to collections of reference amino acid sequences, and automatic annotation using DRAM. Additionally, in some instances, we used phylogenetic trees to confirm functions and identification of motifs determining specific functions (see Materials and methods for details). Solid, partial, and open squares indicate the presence, incompleteness, and absence of gene clusters, respectively. Numbers inside the squares indicate the number of identified amino acid sequences. Abbreviations: PmoCAB: particulate methane monooxygenase (EC: 1.14.18.3); PxmABC: sequence-divergent particulate methane monooxygenase (EC: 1.14.18.3); MmoXYBZ(D)CG: soluble methane monooxygenase (EC: 1.14.13.25); MxaF: calcium-dependent methanol dehydrogenase (subunit 1; EC: 1.1.2.7); XoxF: lanthanide-dependent methanol dehydrogenase (EC: 1.1.2.10); NqrA-F: Na^+^-transporting NADH:ubiquinone oxidoreductase (EC: 7.2.1.1); NuoA-N: NADH:quinone dehydrogenase (EC: 7.1.1.2); CydAB(X): cytochrome bd(-I) (EC: 7.1.1.7); aa3: aa3-type cytochrome c oxidase with the adjacent hemerythrin (EC: 7.1.1.9); caa3: cytochrome c oxidase fused subunit I+III (characteristic for caa3-type cytochrome c oxidase) (EC: 7.1.1.9); Nif and Rsx: molybdenum-dependent nitrogenase complex (>10 genes found, EC: 1.18.6.1) and H^+^/Na^+^-translocating ferredoxin:NAD^+^ oxidoreductase (RsxABCDGE, EC: 7.1.1.11, 7.2.1.2); NarGHI: nitrate reductase (EC: 1.7.5.1, 1.7.99.-); CynS: cyanate lyase (EC: 4.2.1.104); UrtABC: urea transporter; UreABC: urease (EC: 3.5.1.5); SoxB: thiosulphohydrolase (EC: 3.12.1.1); SoxCD: S-disulphanyl-L-cysteine oxidoreductase (EC: 1.8.2.6); Sqr: sulphide:quinone oxidoreductase (EC: 1.8.5.4); BGC: biosynthetic gene cluster. Genomes from our recently cultured strains are shown in **bold**.

**Figure 3.**
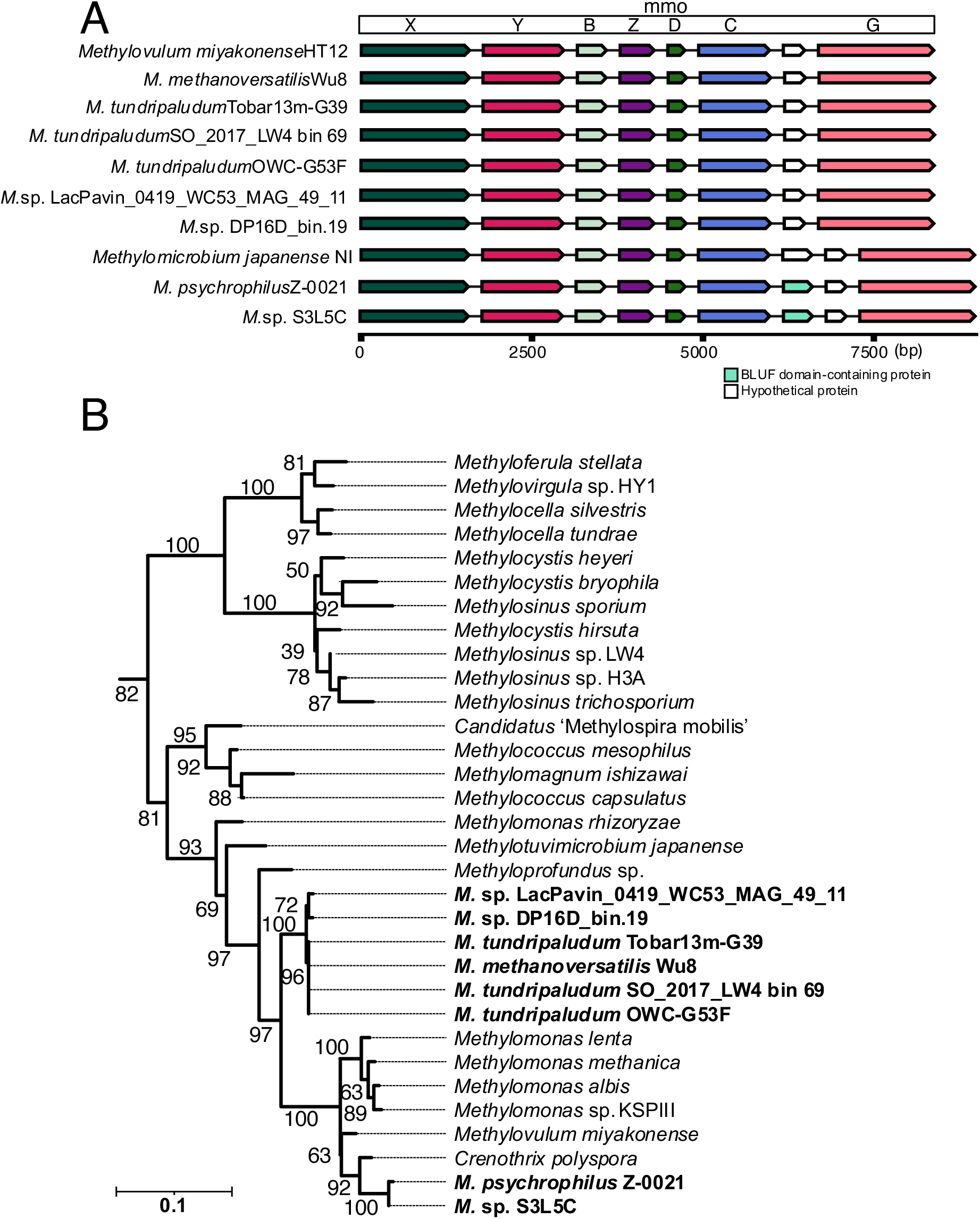
Genomic organisation of the soluble CH_4_ monooxygenase operon and phylogeny of the MmoX subunit among methanotrophs. (A) Gene map of the sMMO operon across representative genomes, illustrating gene order and conservation. Gene lengths are calculated as the raw difference between end and start coordinates (in base pairs, bp), with small gaps added for visualisation; the x-axis is therefore shown in bp. (B) Phylogenetic tree of the MmoX subunit from investigated *Methylobacter* spp. in the context of other methanotrophs from diverse taxonomic groups, showing taxonomic clustering patterns. Branch bootstrap support is indicated by numbers next to the nodes. The scale bar indicated 0.1 amino acid substitutions per site.

Although the presence of sMMO has been reported for the genomes of psychrophilic *M. psychrophilus* Z-0021 (Rissanen *et al*. 2022) and in *M*. sp. S3L5C (Khanongnuch *et al*. 2022), earlier works had failed to identify sMMO in *M. psychrophilus* (Omelchenko *et al*., 1996; Trotsenko and Khmelenina, 2005). The presence of sMMO is rarely associated with *Methylobacter* spp. (Bowman *et al*. 1993; Smith, Costello and Lidstrom 1997; Bowman 2014), or in fact, with the majority of the *Methylococcales* (Semrau 2011; Dedysh and Knief 2018). However, in our survey of >1,000 genomes of *Methylococcales*, MmoX was found in ∼15%, belonging to *Methylococcus*, *Crenothrix*, *Methylomonas, Methyloprofundus, Methylovulum, Methylomagnum,* ‘*Ca*. Methylocalor cossyra’, ‘*Ca.* Methylospira mobilis’, and many uncultured *Methylococcales* (Table S2).

Interestingly, all four strains belonging to the cluster with *M. methanoversatilis* encoded all three CH_4_ monooxygenase gene clusters: pMMO, pXMO, and sMMO (Fig. 2). Although uncommon, the occurrence of all three MMOs has been reported for other methanotrophs, including *Methylocystis hirsuta* CSC1, *Methylocystis bryophila* S285, and *Methylosinus* sp. R-45379 (Han, Dedysh and Liesack 2018; Oshkin *et al*. 2020). Despite its rare occurrence, the presence of all three MMO forms might be an ancestral trait, as all three MMOs have been inferred to be present in extant ancestral proteobacterial methanotrophs (Osborne and Haritos 2018). Likely, this repertoire confers broader metabolic flexibility for energy conservation from CH_4_ oxidation by providing different substrate affinities and specificities (Sullivan, Dickinson, and Chase 1998; Baani and Liesack 2008).

The metabolic flexibility of *Methylobacter* spp. is also evident in their genomic repertoire for the second step of CH_4_ oxidation— the oxidation of methanol to formaldehyde. The majority of analysed genomes encoded for calcium-dependent and lanthanide-dependent methanol dehydrogenase, *mxaFI-MDH* (EC: 1.1.2.7) and *xoxF-MDH* (EC: 1.1.2.10), respectively. The latter contains only one subunit, which was found in almost all analysed *Methylobacter* genomes as a single-copy gene (84% of genomes) or with two copies (11% of genomes) (Fig. 2, Table S3). Xox-MDH can act as a primary methanol dehydrogenase (Chu and Lidstrom 2016) and has been speculated to be more efficient and potentially more metabolically versatile than the calcium-dependent form, for instance, by not only being able to oxidise methanol to formaldehyde but also to formate (Keltjens *et al*. 2014).

### Versatile electron transport chain

Our comparative genomics analyses revealed several variations of electron transport chain components in *Methylobacter*, which are likely adaptations to diverse habitats and provide the capacity to withstand fluctuating environmental conditions. Firstly, we identified redox-driven Na^+^-transporting NADH:ubiquinone oxidoreductase (EC: 7.2.1.1; NQR encoded by *nqrA-F*), which acts as a complex I, in almost all analysed *Methylobacter* spp. (Fig. 2, Table S3). Several genomes in clade I, including those of marine species, encoded two NQRs. Another type of complex I, NADH:quinone dehydrogenase (EC: 7.1.1.2, NUO encoded by *nuoA-N*) was present in the majority of the genomes, but lacking in many *Methylobacter* genomes and MAGs, including the four strains of *M. methanoversatilis* (Fig. 2, Table S3). Both forms of complex I transfer an electron from NADH to quinone, but they pump different cargo to the periplasm: either two H^+^ protons (NUO) or one Na^+^ ion (NQR); however, it has been shown that NQR can also translocate H^+^ protons (Raba *et al*. 2018). Moreover, these complexes differ in their energy conservation capacity, with NQR being likely more energy-efficient compared to NUO (Hreha *et al*. 2021). A fine-tuned differential expression of these two forms has been observed in *M. tundripaludum* SV96, with the NQR dominating expression at temperatures below 15°C (Tveit *et al*. 2023). Additionally, it is likely that the expression of NQR/NUO may be regulated by other factors, such as stoichiometry of Na^+^/e^−^ and H^+^/e^−^, redox, and low O_2_ levels and substrate availability, as indicated for other bacteria (Bogachev, Murtazina and Skulachev 1997; Spero *et al*. 2015; Ito *et al*. 2020; Kaila and Wikström 2021). Certainly, the possibility to choose between them allows those *Methylobacter* spp. that possess both forms to make optimal use of resources for maintaining and building membrane potential. Sole reliance on NQR, on the other hand, may reflect adaptation to elevated salt concentration and/or be necessary for anaerobic metabolism (Buckel *et al*. 2025).

Secondly, we detected up to two high-affinity terminal oxidases of the *bd(-I)*-type (EC: 7.1.1.7; Fig. 2) in the vast majority of investigated *Methylobacter* genomes. These cytochrome *bd* quinol oxidases consist either of two or three subunits (genes *cydAB(X)*). They have been found to enable aerobic respiration under hypoxia (<50 nM O_2_) and withstand nitrosative and oxidative stress better than other types of terminal oxidases (Giuffrè *et al*. 2014). Next to these *bd*-type oxidases, most analysed *Methylobacter spp.* contained other terminal oxidases of the heme-copper A and C types in their genomes (Fig. 2, Table S3). In all genomes containing the *aa3*-type terminal oxidase, which has a low affinity for O_2_ (Berg *et al*. 2022), we found a gene coding for hemerythrin adjacently located, as seen in other gammaproteobacterial methanotrophs (Rahalkar and Bahulikar 2018; Nariya and Kalyuzhnaya 2020). Hemerythrins act as high-affinity O_2_-sensors and carriers whose expression in methanotrophs has been linked to hypoxia (Kalyuzhnaya *et al*. 2013; Rahalkar and Bahulikar 2018; Nariya and Kalyuzhnaya 2020; Weiblen, Sauvageau and Stein 2024) and to high copper concentrations with a pMMO-enhancing function (Kao *et al*. 2008; Chen *et al*. 2012). Since the transcription of these proximally located genes has been proven to be connected to enhanced O_2_-dependent respiration in *Methylomicrobium alcaliphilum* in O_2_-limited conditions (Nariya and Kalyuzhnaya 2020), it is possible that the exact mechanism can occur in other *Methylobacter* spp.

Finally, we identified the presence of Na⁺-translocating ATP synthases next to the H^+^-translocating ones in most analysed genomes (Fig. 2, Table S3). By comparing the amino acid sequences of the subunit c, encoded by *atpE*, from the Na⁺-translocating ATP synthases, we found subunits with one- and two-carboxylate ion coupling motifs (Fig. S5B). The two-carboxylate form has been shown to enable the enzyme to couple Na^+^ transport to ATP synthesis only when Na^+^ is in excess over H^+^ in the environment (Schulz *et al*. 2013; Leone *et al*. 2015). This diversity in components of the respiratory chain, together with the presence of diverse dissimilatory protein complexes (see below), most likely allows *Methylobacter* spp. to employ flexible mechanisms suited for thriving in different, often fluctuating, environmental conditions.

### Potential for using other sources of energy than CH_4_

We identified genes involved in partial denitrification in the majority of the analysed clade II genomes and MAGs. Many of the investigated *Methylobacter* spp. have the potential to reduce nitrate to nitrite (e.g., with the nitrate reductase (EC: 1.7.5.1, 1.7.99.-)), nitrite to nitric oxide (e.g., with the nitrite reductase (NO-forming) (EC: 1.7.2.1)), and nitric oxide to nitrous oxide (e.g., cytochrome c coupled nitric oxide reductase (EC:1.7.2.5)) (Fig. 2, Table S3). However, we did not detect nitrous oxide reductase encoded by *nosZ* reported for some acidophilic methanotrophs belonging to *Alphaproteobacteria* and *Verrucomicrobiia* (Awala *et al*. 2024). Indeed, axenic cultures of *Methylobacter* sp. YRD-M1 growing in O_2_-limiting conditions produced N_2_O or NO (Hao *et al*. 2022).

According to our analyses, all *Methylobacter* genomes contained [NiFe]-hydrogenases of the 1d, 1e, 2c, and 3d groups in various combinations (Fig. 2, Table S3). The groups 1d and 1e are known to be involved in respiratory hydrogen uptake liberating electrons for aerobic and anaerobic respiration (1d) or sulphur respiration (1e), whereas hydrogenases of the group 2c and 3d on the other hand are cytosolic and not known to be coupled to energy conservation (Vignais, Billoud and Meyer 2001; Greening *et al*. 2016; Søndergaard, Pedersen and Greening 2016). Only the hydrogenase of the 3d group was present in all analysed *Methylobacter* genomes and MAGs. They most likely act as redox valves, either providing NADH as a reductant for carbon fixation or catalysing the fermentative production of hydrogen (Vignais, Billoud and Meyer 2001; Søndergaard, Pedersen and Greening 2016). Despite this apparent diversity of hydrogenases present in *Methylobacter* genomes and MAGs, to our knowledge, there is no evidence on the metabolic use of hydrogen from axenically cultured *Methylobacter* spp. This contrasts with the proven hydrogen utilisation by alphaproteobacterial (Hakobyan and Liesack 2020; Hakobyan *et al*. 2020), verrucomicrobial (Carere *et al*. 2019), and gammaproteobacterial methanotrophs with the Calvin-Benson-Bassham cycle (Stanley and Dalton 1982; Liu *et al*. 2024). However, anaerobic laboratory incubations of complex microbial communities from Arctic lake sediments revealed the expression of *Methylobacter* group 1d hydrogenases, suggesting that hydrogen may provide energy for CH_4_ oxidation by *Methylobacter* spp. under O_2_-depleted conditions (He *et al.,* 2022).

Our analyses revealed that sulphide:quinone oxidoreductase (*sqr*; EC: 1.8.5.4) was universally present in all 44 *Methylobacter* genomes and MAGs in one or more copies (Fig. 2, Table S3). A recent study showed that at low CH_4_ concentrations, *Methylobacter* sp. S3L5C used dissolved organic matter and reduced sulphur compounds, such as sulphide, acting as electron donors, while maintaining elevated expression of *sqr* (Rissanen *et al*. 2025). In seven *Methylobacter* spp. from clade I, we additionally detected thiosulphohydrolase (SoxB; EC: 3.12.1.1) along with S-disulphanyl-L-cysteine oxidoreductase (SoxCD; EC: 1.8.2.6), which are components of the periplasmic thiosulphate-oxidising complex catalysing thiosulphate oxidation to sulphate (Fig. 2, Table S3). Therefore, it seems likely that *Methylobacter* spp. are capable of using reduced sulphur compounds as electron donors for growth.

These different types of genes may play important roles in dissimilatory metabolisms; however, especially the hydrogen- and sulphur-related genes are understudied in *Methylobacter* spp. (Fig. 4), and largely in methanotrophs in general. Therefore, it is unclear how much they contribute to energy generation relative to CH_4_ oxidation, and under what conditions these metabolisms could be relevant to the cells.

**Figure 4.**
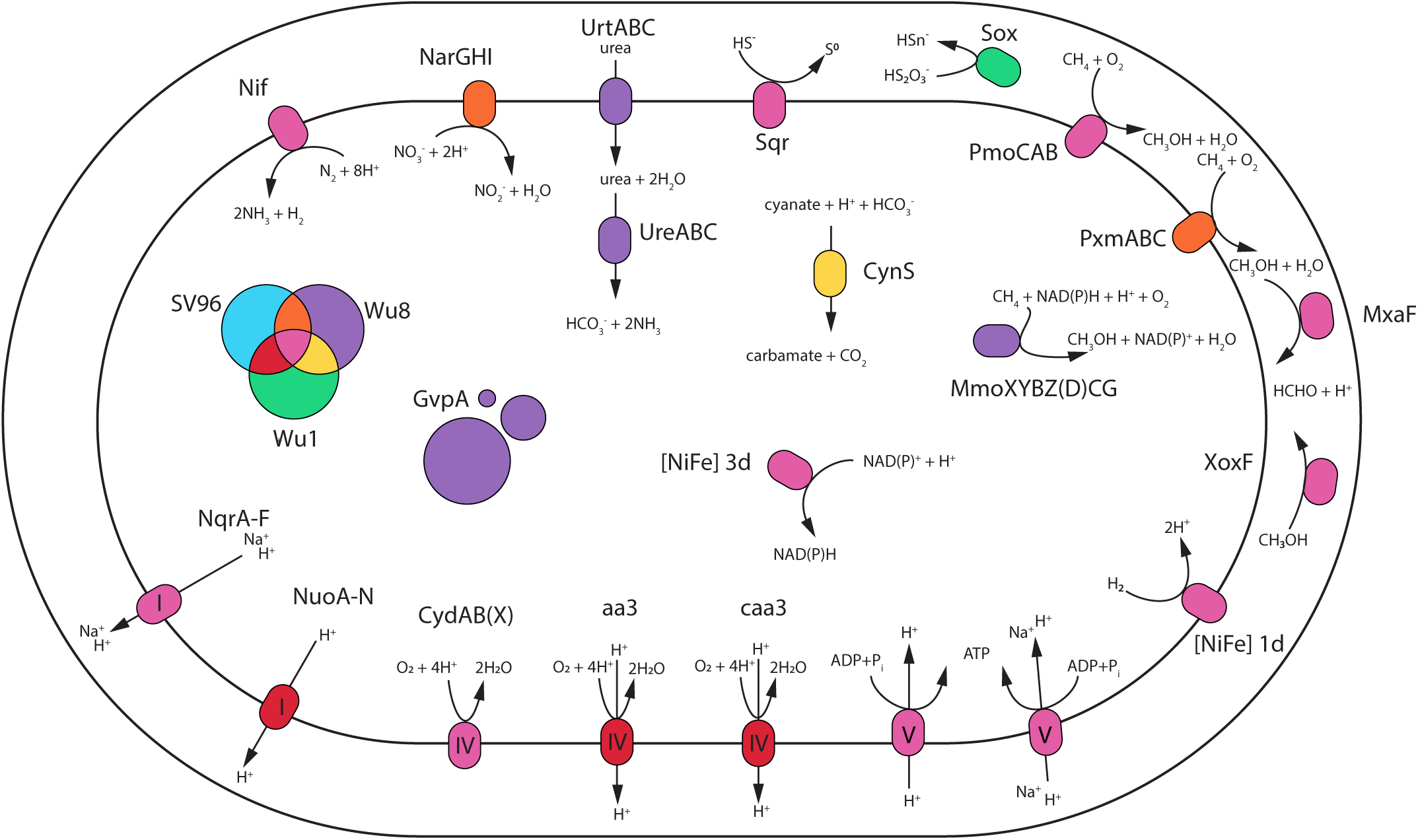
Visual depiction of some of the proteins and protein complexes in three *Methylobacter* spp.: *M. tundripaludum* SV96, *M. methanoversatilis* Wu8 and *M. spei* Wu1. Abbreviations: PmoCAB: particulate methane monooxygenase (EC: 1.14.18.3); PxmABC: sequence-divergent particulate methane monooxygenase (EC: 1.14.18.3); MmoXYBZ(D)CG: soluble methane monooxygenase (EC: 1.14.13.25); MxaF: calcium-dependent methanol dehydrogenase (subunit 1; EC: 1.1.2.7); XoxF: lanthanide-dependent methanol dehydrogenase (EC: 1.1.2.10); NqrA-F: Na^+^-transporting NADH:ubiquinone oxidoreductase (EC: 7.2.1.1); NuoA-N: NADH:quinone dehydrogenase (EC: 7.1.1.2); CydAB(X): cytochrome bd(-I) (EC: 7.1.1.7); aa3: aa3-type cytochrome c oxidase (EC: 7.1.1.9); caa3: cytochrome c oxidase fused subunit I+III (characteristic for caa3-type cytochrome c oxidase) (EC: 7.1.1.9); Nif: molybdenum-dependent nitrogenase complex (>10 genes found, EC: 1.18.6.1); NarGHI: nitrate reductase (EC: 1.7.5.1, 1.7.99.-); CynS: cyanate lyase (EC: 4.2.1.104); UrtABC: urea transporter; UreABC: urease (EC: 3.5.1.5); Sox: potential for oxidising sulphur compounds, such as thiosulphate.

### Diverse genomic potential for nitrogen fixation and assimilation

*Methylobacter* spp. are known for their diverse roles in the nitrogen cycle (Bowman *et al*. 1993; Bowman 2014; Khatri *et al*. 2020). We identified at least 10 genes for molybdenum-dependent nitrogenase (Nif; Fig. 2, Table S3). The presence of Nif coincided with the presence of a gene cluster coding for H^+^/Na^+^-translocating ferredoxin:NAD^+^ oxidoreductase (*rsxABCDGE*, EC: 7.1.1.11, 7.2.1.2), which is architecturally similar to NQR, or Rnf gene cluster found in anaerobic organisms (Koo *et al*. 2003). Rsx transfers electrons to a transcriptional regulator involved in responding to oxidative and nitrosative stress—SoxR, which is reduced during aerobic growth (Ding and Demple 1997; Koo *et al*. 2003). Aerobic dinitrogen fixation has been demonstrated for the Nif and Rsx-containing ‘*Ca.* M. svalbardensis’ (Patil *et al*. 2024), and it is therefore probable that the other *Methylobacter* spp. with the same gene clusters are capable of this metabolism.

*Methylobacter* spp. isolated from various environments can assimilate both nitrate (e.g., with *nasAB*) and ammonia (e.g., with *glnA*, *gdhA*) as they can grow in both nitrate- and ammonia-amended salt media (Bowman *et al*. 1993; Bowman 2014). Additionally, we found that more than half of the analysed species additionally held the potential for assimilation of (reduced) forms of dissolved organic nitrogen, such as cyanate and/or urea (Fig. 2, Table S3). Urea has been identified as a source of nitrogen for several methanotrophs (de la Torre *et al*. 2015; Nguyen *et al*. 2017; Wang *et al*. 2024), and it stimulates the growth of *Methylobacter*-like species (Zheng *et al*. 2014). ‘*Ca*. Methylobacter coli’ strain BlB1 has been shown to grow on urea as a source of nitrogen (Khatri *et al*. 2021); however, in our analyses, we have identified neither urease nor urea-transporting genes (Fig. 2). Moreover, urease may play additional roles, for instance, in the adjustment of internal or external pH (Scott *et al*. 1998; Stingl, Altendorf and Bakker 2002). For other organisms, it has been demonstrated that cyanate serves as a nitrogen source, facilitating growth (Guilloton and Karst 1987; Wood *et al*. 1998). Clearly, however, organic nitrogen uptake by *Methylobacter* spp., and perhaps by all methanotrophs, remains functionally unexplored and understudied.

### Unusual genomic adaptations

Unexpectedly, we found a multigene cluster encoding gas vesicle proteins in all four members of *M. methanoversatilis*, as well as the two psychrophilic species *M. psychrophilus* Z-0021 and *M.* sp. S3L5C, and a groundwater MAG that belonged to *M.* sp. C13 (Fig. 2, Table S3). Typically, the gas protein vesicles gene clusters contain 8–14 genes, and produce a vesicle protein monolayer formed by the gas vesicle protein A (encoded by *gvpA*). The presence of this gene cluster suggests a planktonic lifestyle, as gas vesicles are typically known for enabling organisms to control their buoyancy in the water column (Walsby 1994; Pfeifer 2012). The gas protein vesicle gene cluster has only been reported for two methanotrophs so far—‘*Ca*. Methylomirabilis limnetica’ known for its planktonic lifestyle (Graf *et al*. 2018), and *M. psychrophilus* Z-0021 isolated from tundra soil (Omelchenko, Vasilyeva and Zavarzin 1993). To place our findings in a wider context for gammaproteobacterial methanotrophs, we surveyed >1,000 genomes and MAGs of *Methylococcales* and detected at least one *gvpA* gene copy in ∼12.5% of them (Table S2). Interestingly, they were present in up to five copies of *gvpA*, primarily in the MAGs of uncultured *Methylococcaceae* (NCBI-sourced taxonomy) from O_2_-stratified freshwater bodies in the Northern Hemisphere sequenced in Buck et al. (2021). As some methanotrophs are known to grow preferentially in defined CH_4_-O_2_ gradients (Bussmann, Rahalkar and Schink 2006; Reim *et al*. 2012; Danilova *et al*. 2016; Beals and Puri 2024), perhaps the ability to regulate buoyancy enables them to position themselves at the water column depths with optimal CH_4_:O_2_ stoichiometry. However, five of the seven *Methylobacter* strains in which we detected *gvpA* originate from terrestrial habitats (Fig. 1, Table S1). Other terrestrial bacteria have been reported to contain *gvp* gene clusters (Van Keulen *et al*. 2005), and the function of these organelles remains little understood. As gas vesicles increase the cell surface-to-volume ratio and could therefore improve gas diffusion and aid in survival under stressful conditions (Walsby 1972, 1994; Pfeifer 2012). Indeed, *M. psychrophilus Z-0021* has been shown to produce gas vesicles increasingly with temperatures ranging from 7 to 20°C, but no gas vesicles were detected when the temperature dropped below 7°C (Omelchenko, Vasilyeva, and Zavarzin, 1993). These findings imply that increased gas vesicle production helps to overcome substrate limitation due to lower CH_4_ and O_2_ solubility in the medium with increasing temperature. Protein vesicles are permeable to many gases, including CH_4_ and O_2_ (Walsby 1969, 1971, 1982), but it is unclear which gas(es) methanotrophs contain in the vesicles. It is tempting to speculate that the accumulated gas(es) could serve as a temporal reservoir of metabolically important gases, such as O_2_ or CH_4_. However, these speculations require appropriate experimental verification.

On average, we identified nearly eight biosynthetic gene clusters (BGCs) per analysed *Methylobacter* genome/MAG, ranging from three in the genomes of organisms from nutrient-poor, pelagic environments to 15 in a MAG from freshwater sediment (Fig. 2). The most attention among *Methylobacter* spp. BGCs was given to tundrenone, a quorum-sensing molecule likely involved in hypoxia stress response (Puri et al., 2018; Yu et al., 2020), and to ectoine, an organic osmoprotectant that, among other functions, increases halotolerance (Reshetnikov et al., 2011). Nevertheless, we could only confirm the complete tundrenone gene cluster (encoded by *tunA-P* and several regulatory genes on both sides of the gene cluster) for *M. tundripaludum* SV96, 21/22, and 31/32 (Fig. 2). Similarly, a complete ectoine BGC was only detected in three genomes of the clade I: *M. marinus* A45, *M*. sp. BBA5.1, and *M*. sp. B4-02 (Fig. 2). Moreover, most genomes/MAGs contained more than one BGC encoding terpenes, a metabolite group involved in communication or interaction between species, that has been shown to be produced by *M. luteus* in the presence of *Pseudomonas mandellii* (Veraart *et al*. 2018). Furthermore, all analysed *Methylobacter* contained at least one BGC for redox-cofactors and aryl polyenes. The latter have recently been reported as common in gammaproteobacterial methanotrophs (Krause *et al*. 2025) and may encode antioxidative pigments (Schöner *et al*. 2016). Finally, we detected an abundance of diverse, yet unexplored, BGC gene clusters encoding for polyketides, which are known to often carry antibiotic and pharmacological properties. Impressively, two BGCs contained polyketide synthetases that were among the longest genes in *M. methanoversatilis*, spanning over 18.5k (NCBI Reference Sequence: WP_331306173.1) and 26k bp (NCBI Reference Sequence: WP_331307029.1). Despite the availability of genomic information and the presence of gene clusters with high interest for applied science, the biotechnological potential of *Methylobacter* spp. has not been fully explored. Genomic data can be used for metabolic modelling (Islam *et al.,* 2020; Wutkowska *et al.,* 2024) to describe the functioning of *Methylobacter* spp. alone and within microbial communities, which may direct future design of biotechnological applications.

### Genome-inferred optimal growth conditions may guide future cultivation efforts

A large diversity of methanotrophs, including most analysed *Methylobacter* spp., remains uncultured, which hampers the investigation of genome-derived hypotheses regarding their metabolism and ecological niche. To assist and possibly direct cultivation efforts, we predicted optimal conditions for growth regarding temperature, salinity, and pH, as well as tolerance to oxygen from the 44 high-quality genomes and MAGs (Fig. S6). Comparing the predictions with experimentally verified optima for those species that are cultured showed that the genome-wide amino acid composition predictions do not identify the exact optima (Fig. S6). However, general temperature and salinity preferences were likely accurately predicted; for example, the predicted temperature optima for known psychrophiles were among the lowest (Fig. S6a), and known halophilic species and species from oligotrophic environments were predicted to fall at opposite ends of the salinity range (Fig. S6c). Contrastingly, the predictions of optimal pH failed to confirm known pH preferences (Fig. S6b) and were associated with large uncertainties. Without further studies, it is challenging to identify the cause of these discrepancies.

Although all *Methylobacter* genomes and MAGs in this study were expectedly tagged as oxygen-tolerant and are classified as aerobic CH_4_ oxidisers, an increasing number of studies reported *Methylobacter* spp. found in O_2_-limited or O_2_-depleted environments (Nercessian *et al*. 2005; Biderre-Petit *et al*. 2011; Graef *et al*. 2011; Reim *et al*. 2012; Hernandez *et al*. 2015; Oshkin *et al*. 2015; Martinez-Cruz *et al*. 2017; Rissanen *et al*. 2018; Singleton *et al*. 2018; Cabrol *et al*. 2020; Mayr *et al*. 2020; van Grinsven *et al*. 2020b, 2020a; Hao *et al*. 2022; He *et al*. 2022; Grégoire, George and Hug 2023; Li *et al*. 2023, 2024, 2025; Gafni *et al*. 2024; Kallistova *et al*. 2024; Reis *et al*. 2024; Schorn *et al*. 2024), and sometimes able to outcompete other methanotrophs (Beck *et al*. 2013; Hernandez *et al*. 2015; Islam *et al*. 2020; Li *et al*. 2025). To date their persistence or continuous activity under O_2_-limited and O_2_-depleted conditions is far from being fully understood (Hernandez *et al*. 2015; Gafni *et al*. 2024; Li *et al*. 2024; Reis *et al*. 2024; Ruff *et al*. 2024), but in the analysed *Methylobacter* genomes and MAGs we found many genes that provide insights into their functioning under O_2_-depleted conditions, such as nitrate reductases, high-affinity terminal oxidases, hemerythrins, hydrogen uptake [NiFe]-hydrogenases, and the inventory for sulphide and thiosulphate oxidation (Fig. 2. Table S3). Evidence for additional genes responding to O_2_-depleted conditions is being continuously reported. For instance, a recent transcriptomics study found that some proteins within the type VI secretion system operon, which are present only in genomes of the clade I, including *M. luteus*, were upregulated in low O_2_ conditions (Beals and Puri, 2024). Attempting to culture and isolate new *Methylobacter* spp. in carefully designed O_2_-depleted conditions, therefore, represents a promising avenue to a better understanding of their spread in the environment.

### Conclusions

Despite apparent environmental importance and widespread occurrence, *Methylobacter* species are still poorly understood in many aspects. Our study shows that the genus harbours more species than currently described, and some of them cannot be distinguished based on 16S rRNA gene sequence comparison. *Methylobacter* species display diverse adaptations to their environments, i.e. diverse combinations of the three different CH_4_ monooxygenases and the potential for several dissimilatory metabolisms. We would also like to encourage testing some of the metagenomic-based hypotheses that we outlined here to move to the post-genomic era with the *Methylobacter* spp.

### Taxonomic considerations

#### Description of *Methylobacter spei*, sp. nov

Etymology: L. fem. n. *spes*, hope; genitive singular *spei*, meaning “of hope”, since the organism was enriched from a fishpond in SE Czech Republic called Hope (Naděje).

A freshwater aerobic chemoorganotroph that oxidises CH_4_ was obtained from organic-rich sediment in an eutrophic fishpond near Hluboká nad Vltavou, called Hope (Naděje). Phylogenetically affiliated with the genus *Methylobacter*, family *Methylomonadaceae*, order *Methylococcales*, phylum *Pseudomonadota*. The genome is 100% complete with 0.36% contamination (CheckM2). It consists of 85 contigs of a total length of 4.4 Mbp and 52% GC content. The coding density is 86.5% with 4,078 total predicted protein sequences and an average gene length of 311.5 bp.

The genome Wu1^Ts^ represented by a MAG (GenBank accession: GCA_036553575.1, GCF_036553575.1), is designated the nomenclatural type for the species, and was recovered from a low-complexity enrichment containing three species.

#### Description of Methylobacter methanoversatilis, sp. nov

Etymology: *methyloversatilis* N.L. neut. n. *methanum*, methane; L. adj. *versatilis*, versatile, adaptable; N.L. adj. *methanoversatilis*, referring to the potential metabolic versatility of the organism in utilizing CH_4_ with three diverse CH_4_ monooxygenases, i.e. pMMO, pXMO and sMMO, encoded in the genomes.

A freshwater aerobic chemoorganotroph that oxidises CH_4_ was obtained from organic-rich sediment of a eutrophic fishpond in the vicinity of Hluboká nad Vltavou called Hope (Naděje). Phylogenetically affiliated with the genus *Methylobacter*, family *Methylomonadaceae*, order *Methylococcales*, phylum *Pseudomonadota*. The proposed species is represented by 4 MAGs: *Methylobacter tundripaludum* OWC-G53F (GenBank accession: GCA_002934365.1 and GCF_002934365.1), *Methylobacter tundripaludum* Tobar13m-G39 (GenBank accession: GCA_021736725.1), *Methylobacter tundripaludum* SO_2017_LW4 bin 69 (GenBank accession: GCA_023227865.1), *Methylobacter* sp. Wu8 (GenBank accession: GCA_036440755.1 and GCF_036440755.1), collected in 4 different geographic localities. The latter was obtained from an axenic culture that is no longer available. None of the genomes are complete/circular, however, they are of high quality and low contamination (Table below). The genome size is 4.03–4.18Mbp, with the G+C content 51–52%, coding density 88.1–88.3%, total predicted coding sequences 3,715–3,788, and average gene length 319.5–327.7bp. The metabolic predictions indicated that the organism encodes for three different CH_4_ monooxygenases, i.e. two particulate (pMMO and pXMO) and soluble (sMMO).

|  | Size | Compl | Cont | CD | protein | avgene len | origin |
| --- | --- | --- | --- | --- | --- | --- | --- |
| GCA_002934365.1 | 4.18 | 99.92 | 0.0 | 88.1% | 3754 | 327.7 | USA |
| GCA_021736725.1 | 4.05 | 99.86 | 0.01 | 88.3% | 3719 | 320.6 | Spain |
| GCA_023227865.1 | 4.03 | 91.74 | 2.29 | 88.2% | 3715 | 319.5 | USA |
| GCA_036440755.1 | 4.16 | 95.92 | 0.0 | 88.1% | 3788 | 322.8 | Czechia |

The genome Wu8^Ts^ represented by a MAG available under the GenBank accession: GCA_036440755.1, is designated nomenclatural type for the species, and was recovered from temperate eutrophic fishpond sediments.

## Data availability

The data were derived from sources in the public domain. 97 *Methylobacter* spp. genomes and MAGs were downloaded from NCBI Genome portal on May 21, 2024, from https://www.ncbi.nlm.nih.gov/datasets/genome/?taxon=429.1067 *Methylococcaceae* genomes and MAGs were downloaded from NCBI Genome portal on August 12, 2024, from https://www.ncbi.nlm.nih.gov/datasets/genome/?taxon=403.

Code generated to analyse the data is deposited at https://github.com/magdawutkowska/methylobacter_comparative_genomics/.

The names of the novel *Methylobacter* species *Methylobacter methanoversatilis*, sp. Nov. and *Methylobacter spei*, sp. nov. have been registered under the SeqCode: https://seqco.de/i:52928 and https://seqco.de/i:52927, respectively.

## CRediT author statement

MW: conceptualization, formal analysis, visualisation, writing - original draft, writing - review & editing; JAN: formal analysis, writing - review & editing; VT: methodology, formal analysis, writing - review & editing; JEN: formal analysis, writing - review & editing; AD: conceptualization, funding acquisition, formal analysis, writing - review & editing.

## Supplementary figures

**Supplementary Figure S1.**
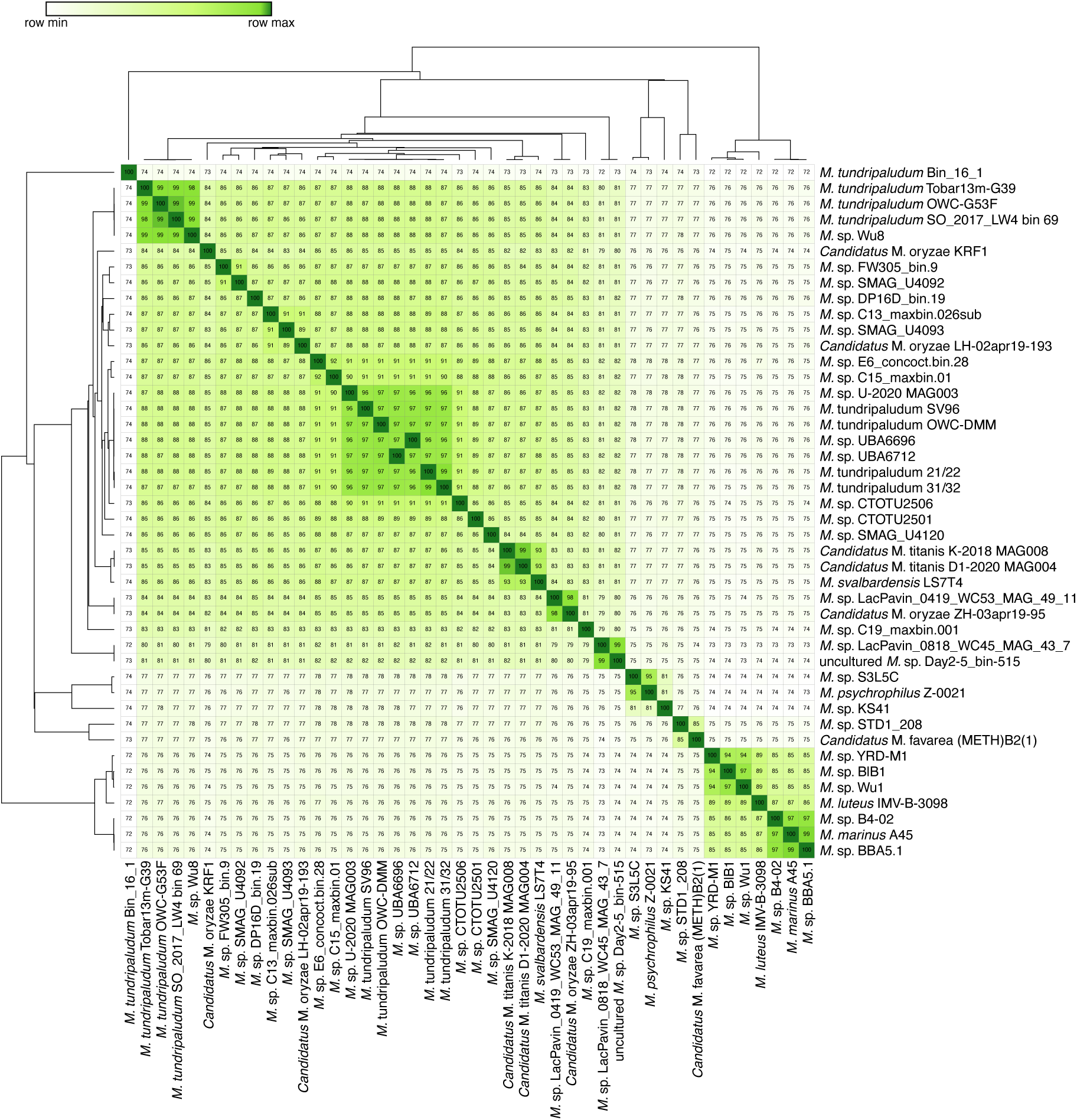
Average amino acid identity (AAI) clustering of the investigated *Methylobacter* spp. genomes and MAGs. The genome AAI similarity matrix was visualised using the Morpheus web-based tool (https://software.broadinstitute.org/morpheus/). Hierarchical clustering was performed using one minus Pearson correlation as the distance metric and average linkage. Both rows and columns were clustered to identify coherent genome groups.

**Supplementary Figure S2.**
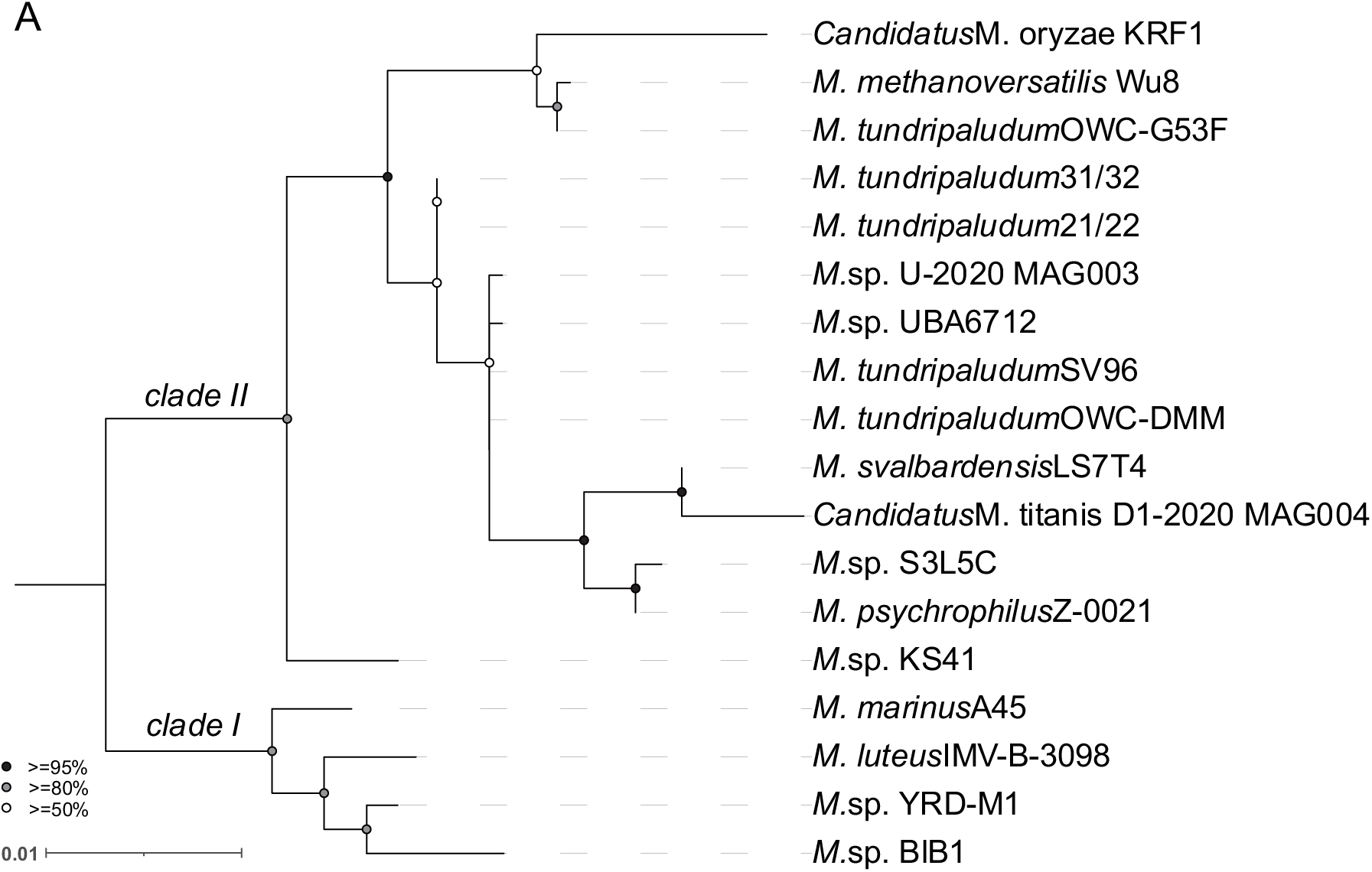
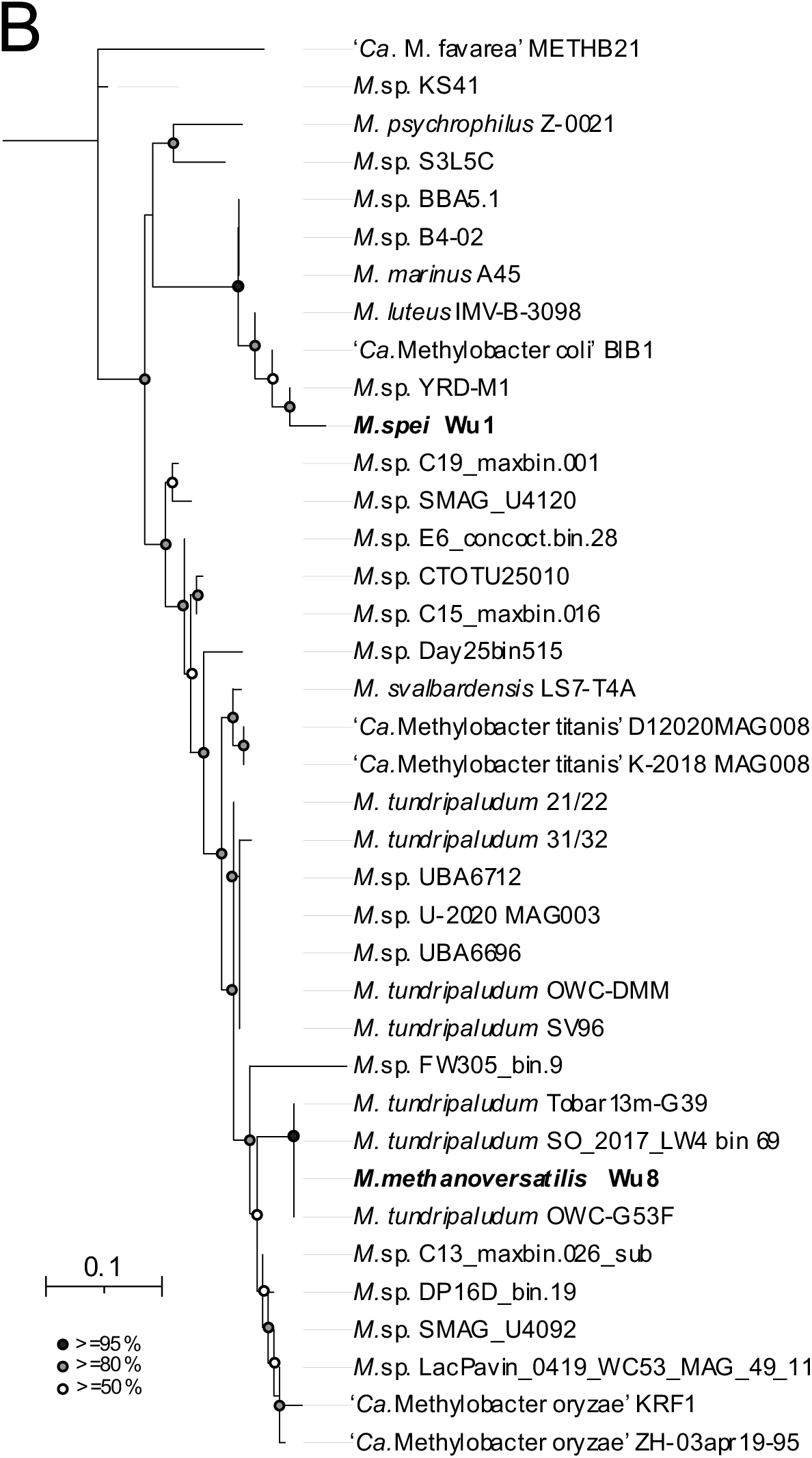
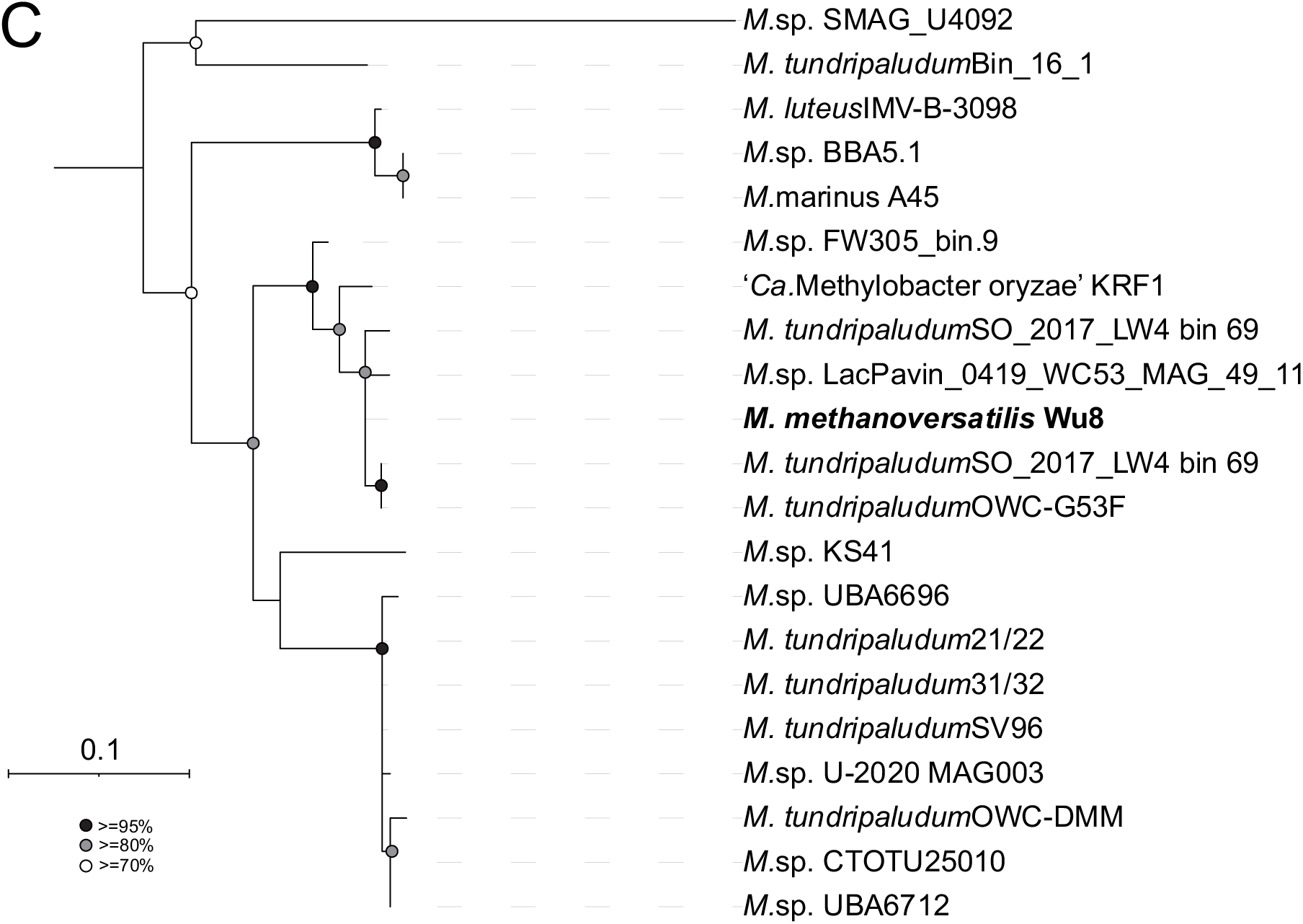
Phylogenetic trees of genes extracted from the investigated *Methylobacter* spp. **Panel A** shows the full-length 16S rRNA gene (>1,500 bp); **panel B**: subunit A of the particulate methane monooxygenases (pMMO), and **panel C**: subunit A of the sequence-divergent particulate methane monooxygenase (pXMO). The 16S rRNA tree was rooted with *Methylomicrobium lacus* LW14 full 16S rRNA nucleotide sequence (NCBI accession number: NR_042712.1). The methane monooxygenase tree was made with all the identified particulate methane monooxygenases, except partial sequences, and was rooted with the AmoA amino acid sequence from *Nitrosomonas europaea* strain ATCC 19718 (UniProtKB/Swiss-Prot: Q04507.2). Bootstrap values are represented by circles in three colours: black (>= 95%), grey (80-95%) and white (50-80%).

**Supplementary Figure S3.**
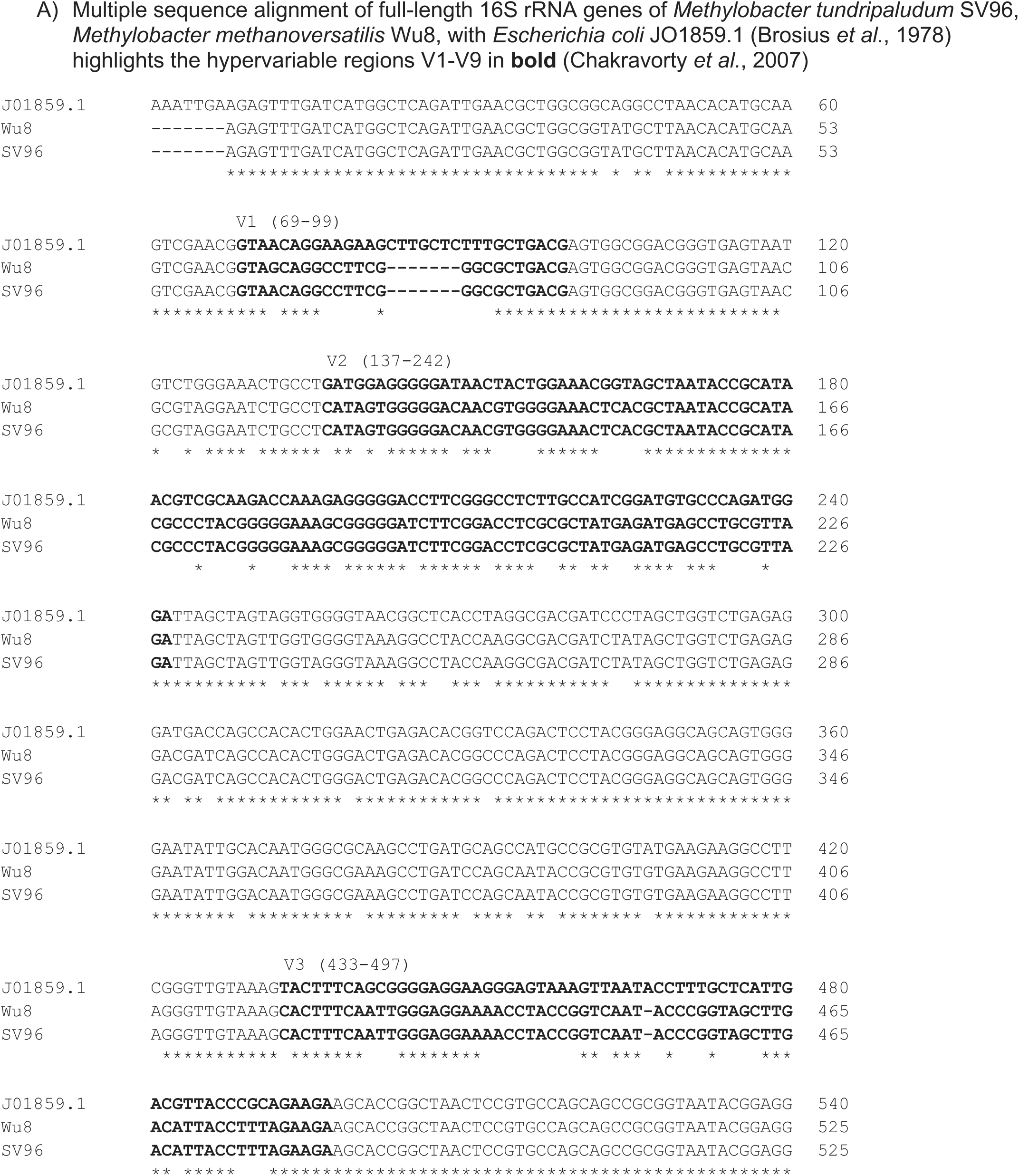
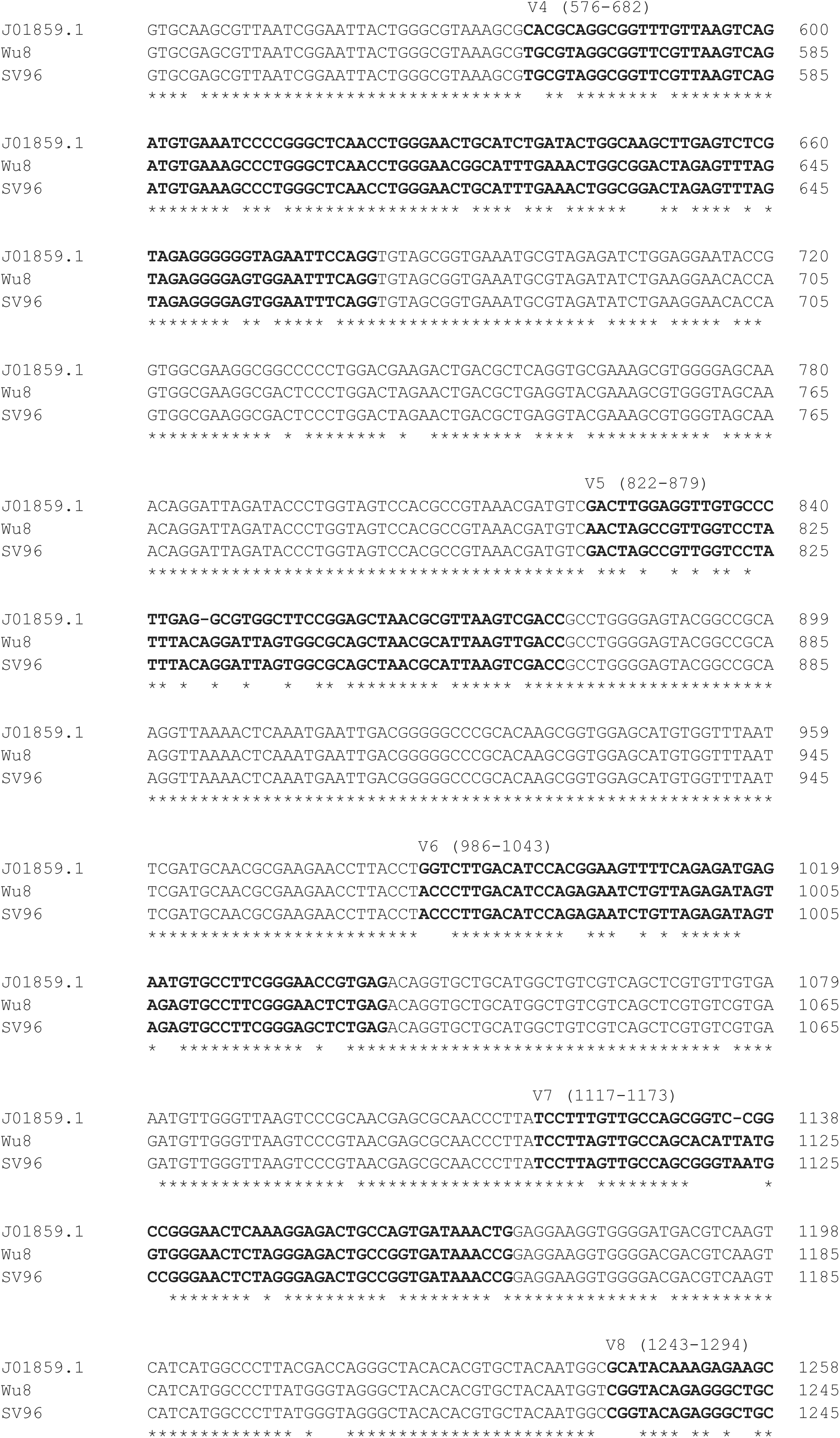
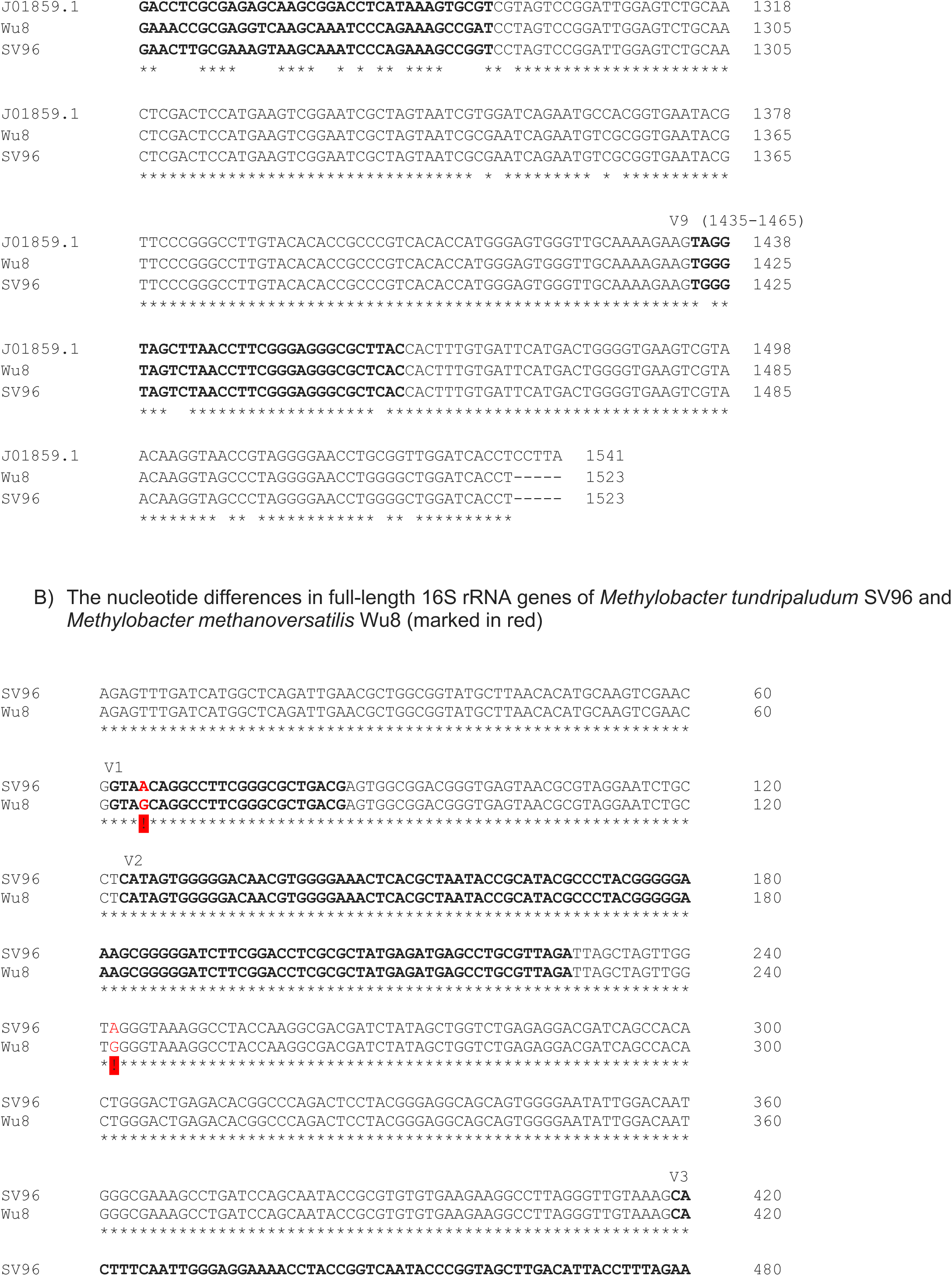
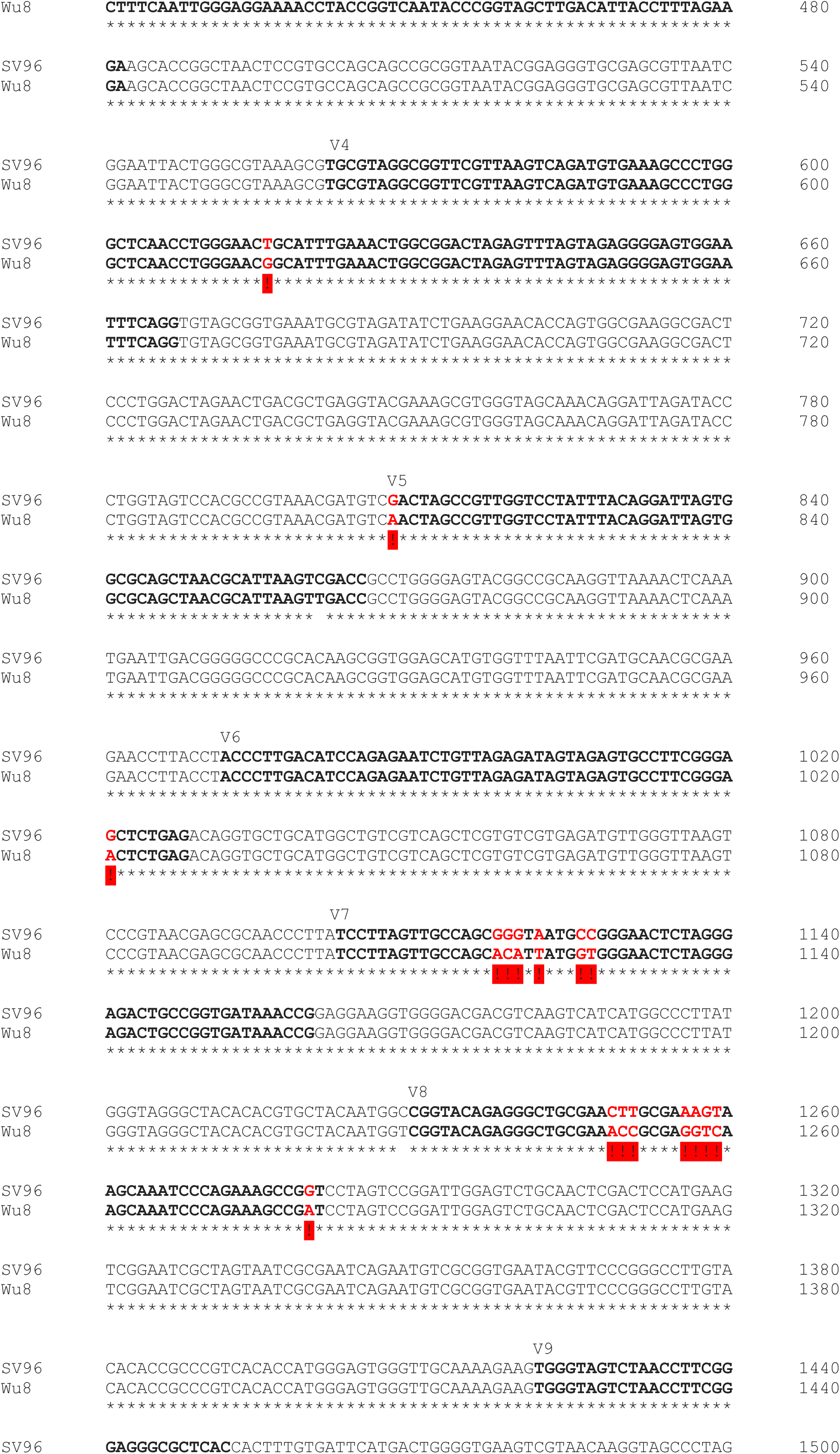
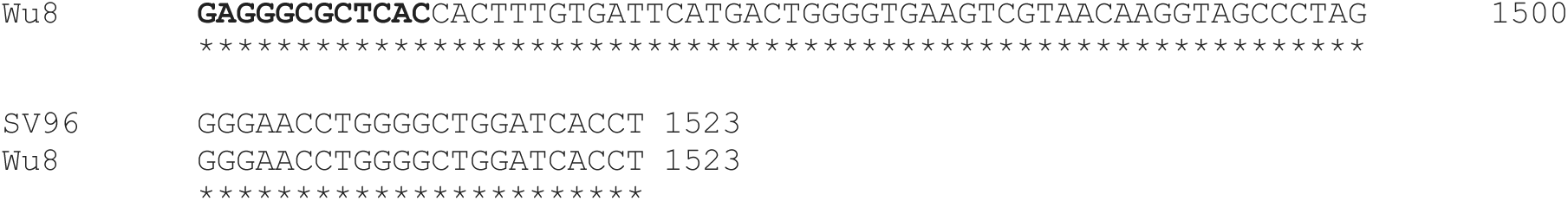
Comparison of 16S rRNA genes in *Methylobacter tundripaludum* SV96 and *Methylobacter methanoversatilis* Wu8. Panel A shows multiple sequence alignment of these two sequences with 16S rRNA of *Escherichia coli* JO1859.1 (Brosius *et al*. 1978), highlighting the hypervariable regions V1-V9 in bold (Chakravorty *et al*. 2007). Panel B shows the differences between the two methanotrophic 16S rRNA sequences in red. A) Multiple sequence alignment of full-length 16S rRNA genes of *Methylobacter tundripaludum* SV96, *Methylobacter methanoversatilis* Wu8, with *Escherichia coli* JO1859.1 (Brosius *et al*., 1978) highlights the hypervariable regions V1-V9 in **bold** (Chakravorty *et al*., 2007)

**Supplementary Figure S4.**
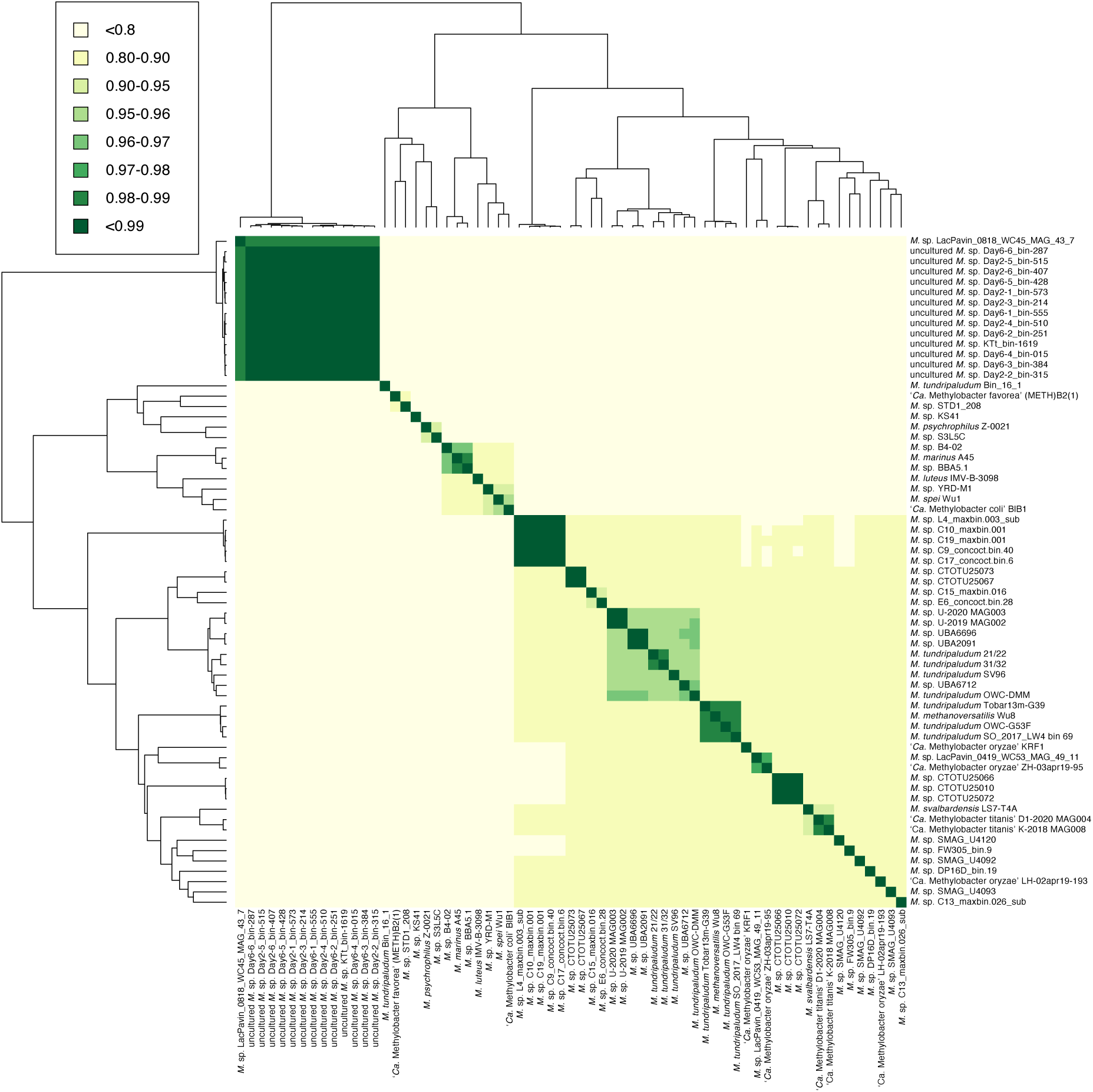
Average nucleotide identity (ANIb) clustering of the investigated *Methylobacter* spp. genomes and MAGs.

**Supplementary Figure S5.**
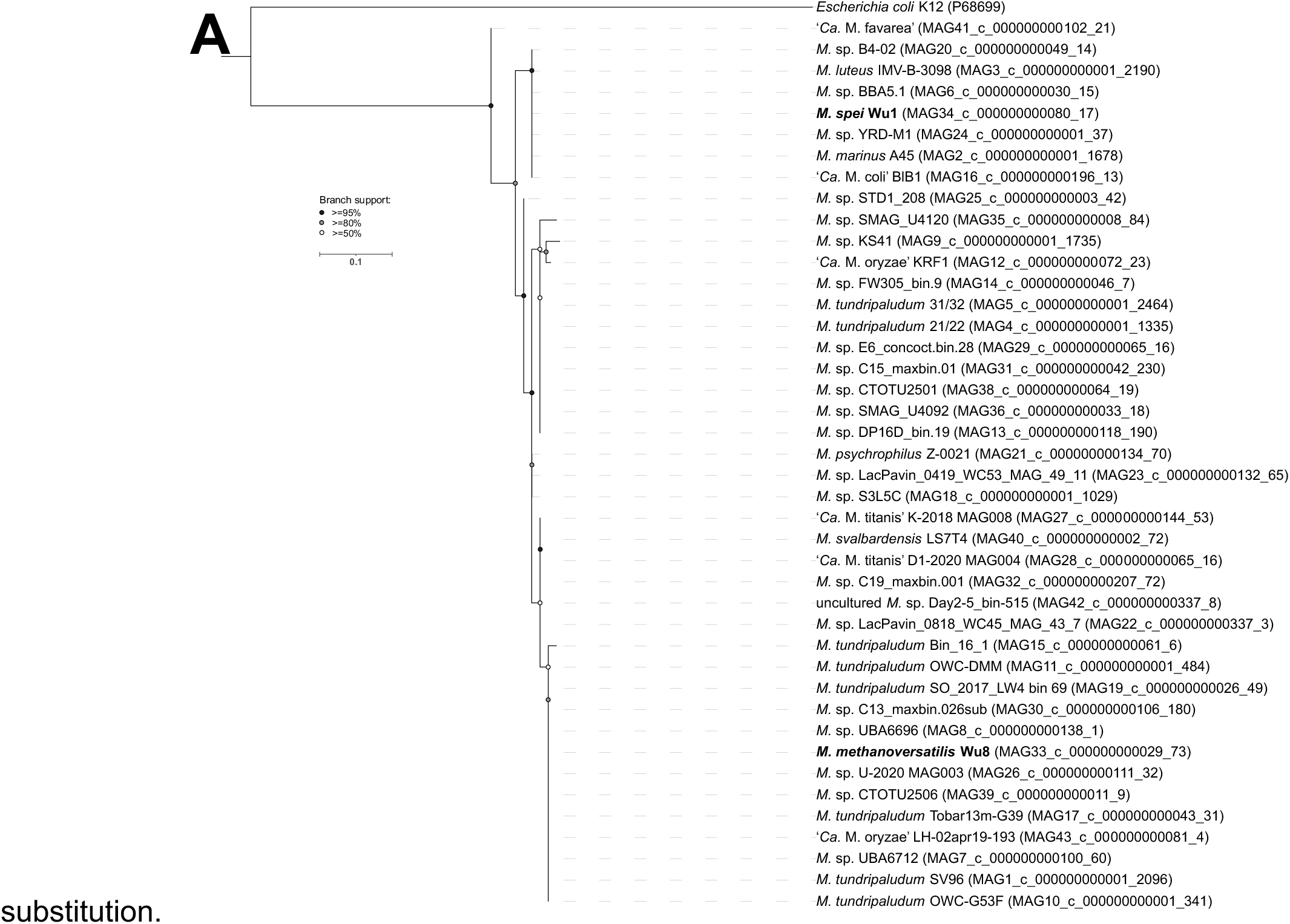
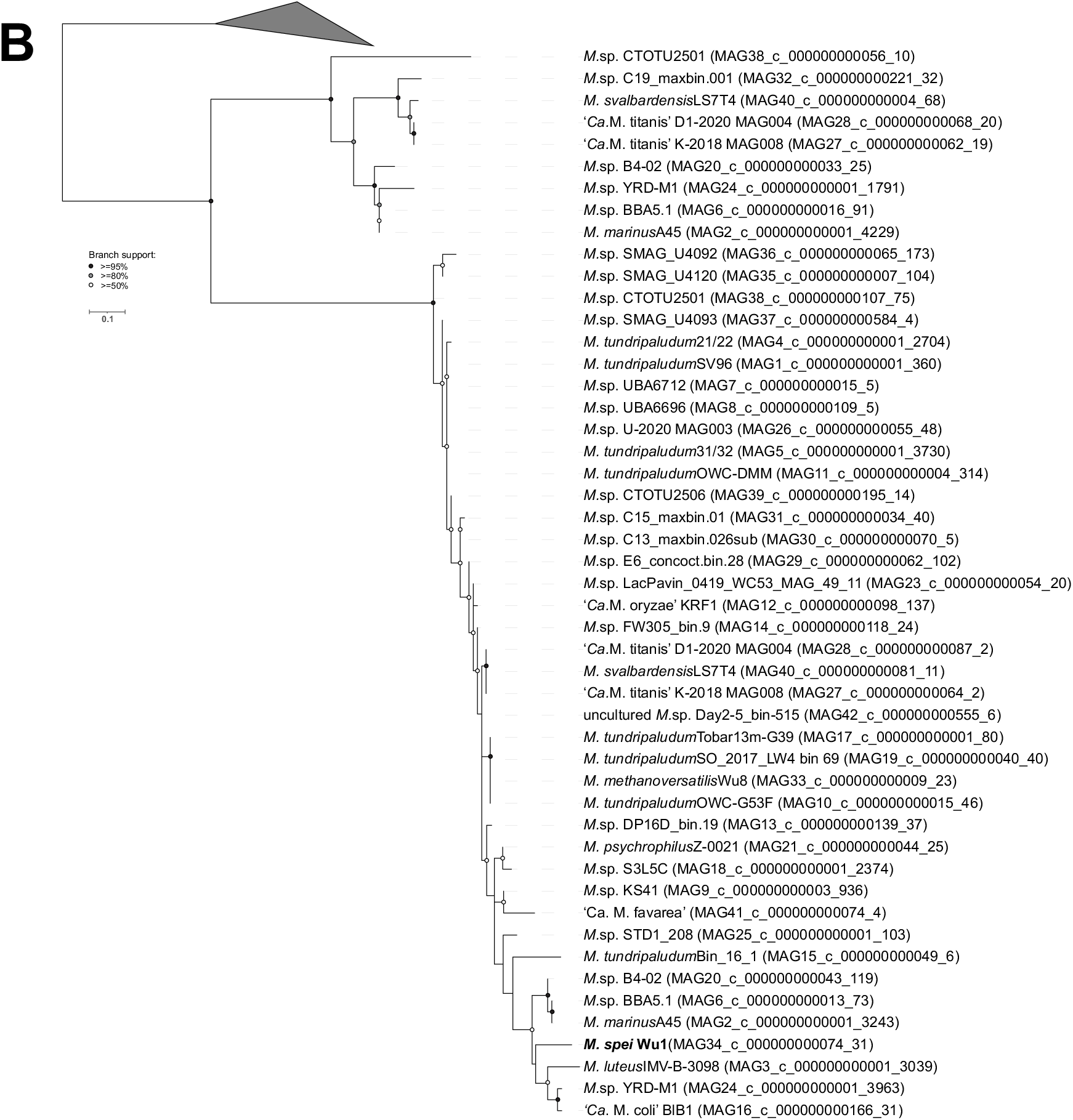
Unrooted phylogenetic trees of ATP synthase subunit 3 (panel A: identified H^+^-ATP synthase AtpE; panel B: identified Na^+^-ATP synthase AtpE). Collapsed clade on panel B contains AtpE sequences that are known for Na^+^-ATP synthase activity. The scale bar indicated 0.1 amino acid substitutions per site. Branch bootstrap support is shown on the nodes as black (≥95%), grey (≥80%), or white (>50%) circles. The scale bar indicates 0.1 amino acid

**Supplementary Figure S6.**
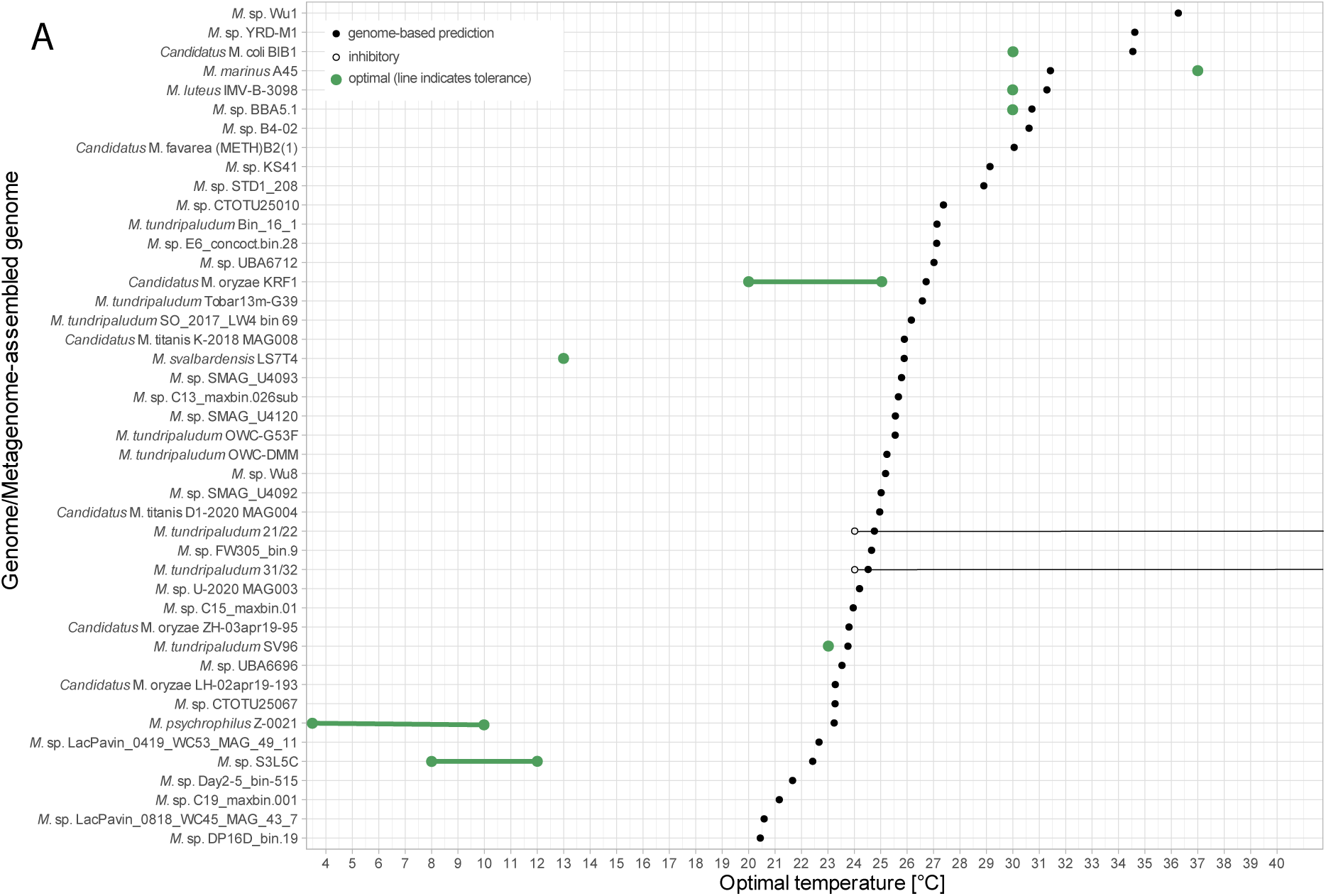
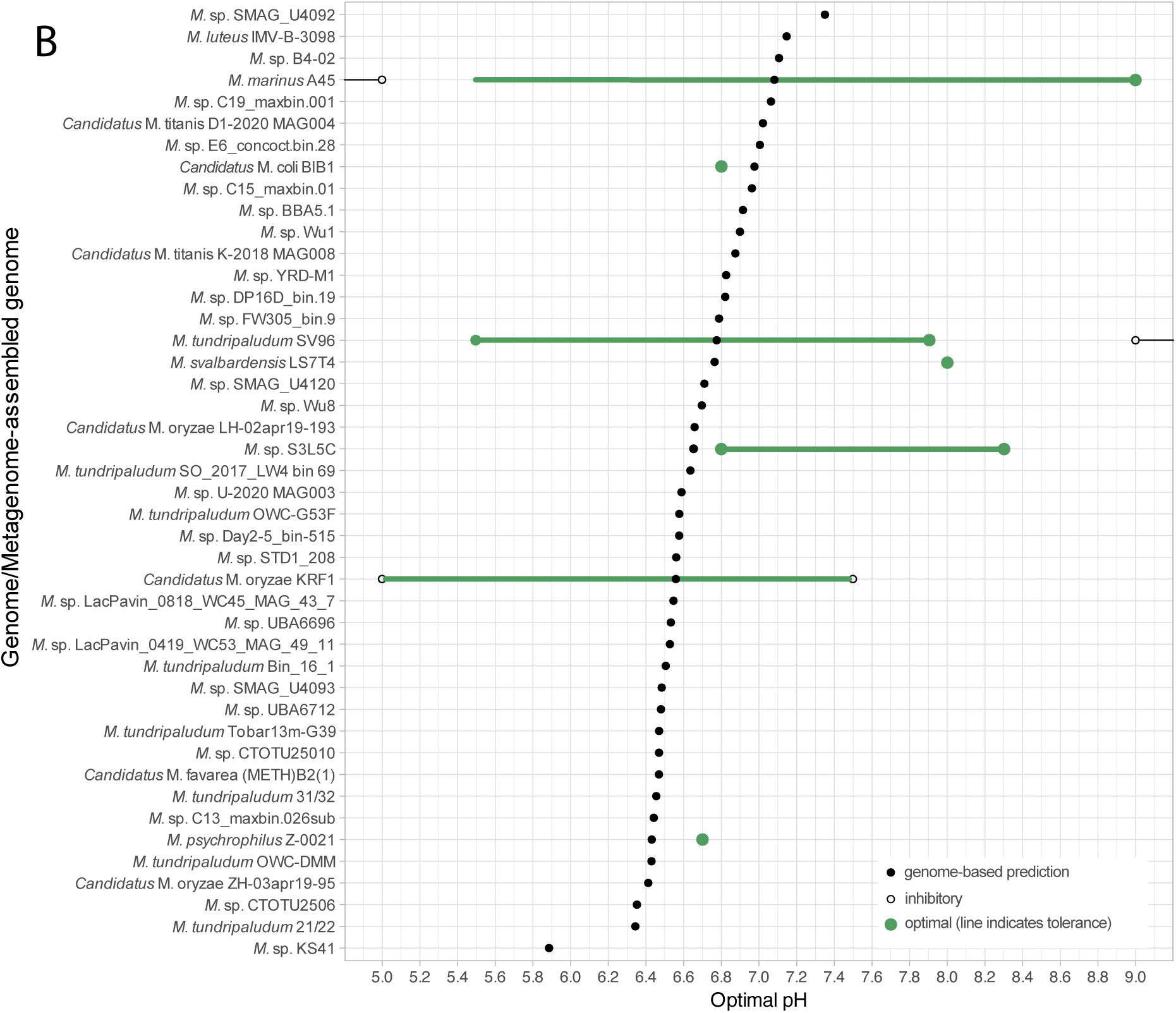
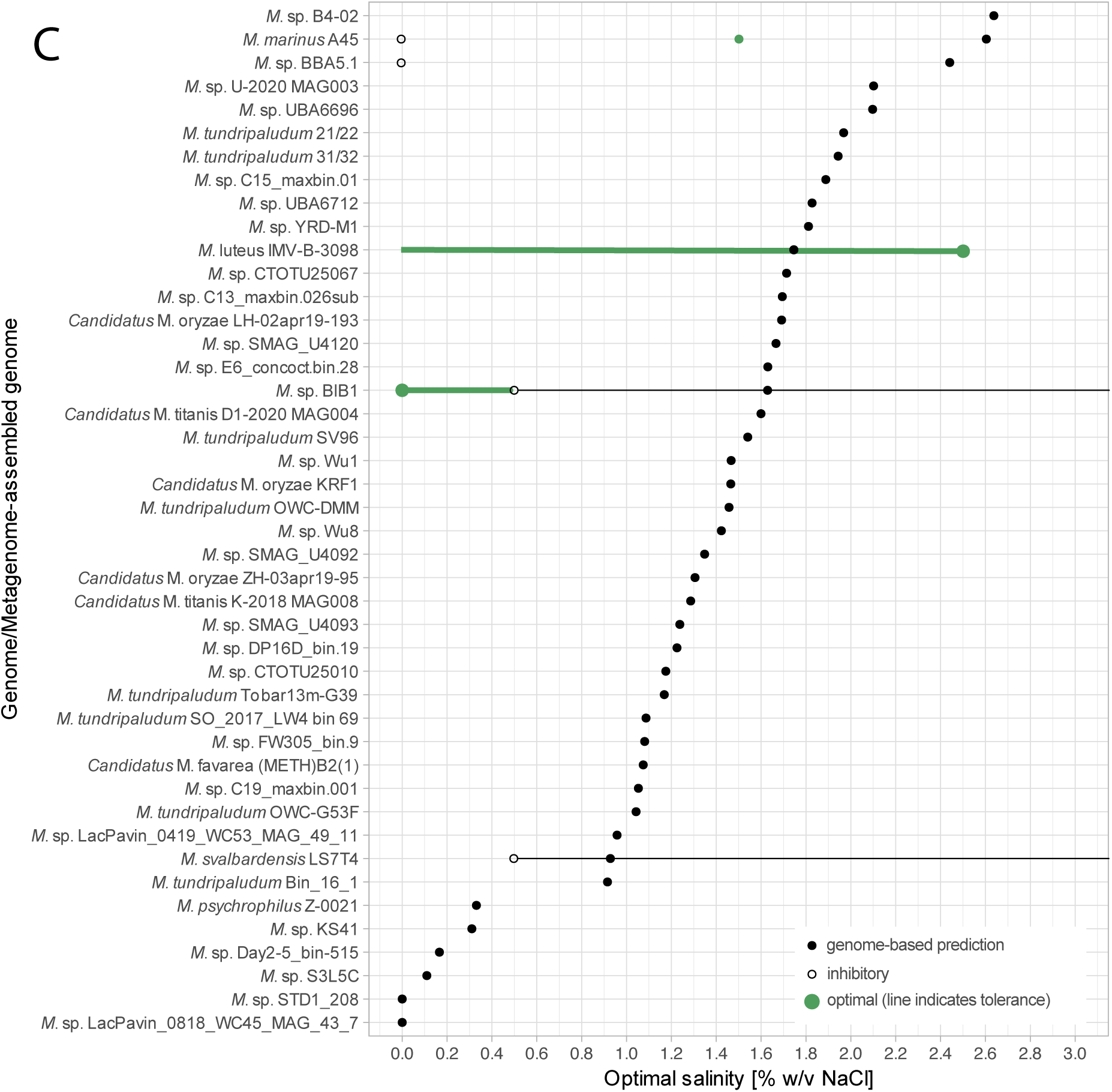
Genome-based predictions of putative optimal temperature (A), pH (B) and salinity (C) values for culturing the investigated *Methylobacter* spp. Green lines indicate verified growth ranges of cultured species. Closed and open circles (with attached thin black lines) indicate predicted optima and experimentally verified values, respectively.

## Supplementary tables

https://github.com/magdawutkowska/methylobacter_comparative_genomics/blob/main/Supplementary_tables.ods

**Supplementary Table S1**

The collection of 97 *Methylobacter* spp. genomes and MAGs deposited in the NCBI Genome database before May 21, 2024. The table contains database identifiers, metadata, and descriptors, such as GenBank assembly accession, accession version, taxonomic assignment from NCBI and GTDB, origin of geographic localities and habitats, completeness and contamination (inferred from CheckM2 analysis), genome/MAG characteristics, and inclusion criteria (columns W and X). Columns W indicate whether the genome passed the high-quality mark (>90% completeness and <5% contamination), whereas column X indicates which genomes/MAGs were included in the high-quality non-redundant dataset included in the analyses based on ANIb clustering. Genomes/MAGs clustered >99% ANIb similarity were labelled as redundant and not included in the dataset (see Fig. S2).

**Supplementary Table S2.**

Copy number of genes encoding for GvpA (the major gas protein vesicle protein) and MmoX (a subunit of soluble methane monooxygenase) in 1,067 genomes and MAGs from NCBI-sourced *Methylococcales*.

**Supplementary Table S3.**

Automatic gene annotations from DRAM.

